# Best organic farming expansion scenarios for pest control: a modeling approach

**DOI:** 10.1101/2022.05.31.494006

**Authors:** Thomas Delattre, Mohamed-Mahmoud Memah, Pierre Franck, Pierre Valsesia, Claire Lavigne

## Abstract

Organic Farming (OF) has been expanding recently in response to growing consumer demand and as a response to environmental concerns. The area under OF is expected to further increase in the future. The effect of OF expansion on pest densities in organic and conventional crops remains difficult to predict because OF expansion impacts Conservation Biological Control (CBC), which depends on the surrounding landscape (i.e. both the crop mosaic and semi-natural habitats). In order to understand and forecast how pests and their biological control may vary during OF expansion, we modeled the effect of spatial changes in farming practices on population dynamics of a pest and its natural enemy. We investigated the impact on pest density and on predator to pest ratio of three contrasted scenarios aiming at 50% organic fields through the progressive conversion of conventional fields. Scenarios were 1) conversion of Isolated conventional fields first (IP), 2) conversion of conventional fields within Groups of conventional fields first (GP), and 3) Random conversion of conventional field (RD). We coupled a neutral spatially explicit landscape model to a predator-prey model to simulate pest dynamics in interaction with natural enemy predators. The three OF expansion scenarios were applied to nine landscape contexts differing in their proportion and fragmentation of semi-natural habitat. We further investigated if the ranking of scenarios was robust to pest control methods in OF fields and pest and predator dispersal abilities.

We found that organic farming expansion affected more predator densities than pest densities for most combinations of landscape contexts and OF expansion scenarios. The impact of OF expansion on final pest and predator densities was also stronger in organic than conventional fields and in landscapes with large proportions of highly fragmented semi-natural habitats. Based on pest densities and the predator to pest ratio, our results suggest that a progressive organic conversion with a focus on isolated conventional fields (scenario IP) could help promote CBC. Careful landscape planning of OF expansion appeared most necessary when pest management was substantially less efficient in organic than in conventional crops, and in landscapes with low proportion of semi-natural habitats.

## Introduction

The intensification of agricultural systems in recent decades has relied on both an increase in field area and a larger dependency on chemical pesticides (Duru et al., 2015; Knapp & van der Heijden, 2018). This process led to profound changes in landscapes with a reduction and fragmentation of semi-natural habitats (Sirami et al., 2019; Tscharntke et al., 2016) and a reduced diversity of the crop mosaic (Tscharntke et al., 2016). This process had demonstrated negative impacts on biodiversity (Perez-Alvarez et al., 2019; Sánchez-Bayo & Wyckhuys, 2019) including on species of interest for agriculture such as pollinators (Goulson Dave et al., 2015; Muth & Leonard, 2019) and pest natural enemies (Sánchez-Bayo & Wyckhuys, 2019; Sirami et al., 2019; Tsutsui et al., 2018). Further, the intensive use of chemical pesticides had negative impacts on human health, and biodiversity (Mózner et al., 2012; Pärn et al., 2012; Sánchez-Bayo & Wyckhuys, 2019). Hence, there is an urgent need to identify alternative farming systems that reduce the negative impacts of intensive agriculture while maintaining yields as much as possible (Colbach et al., 2020; Smith et al., 2020). Organic Farming (OF) is one of these possible alternative systems. The multi-performance of OF recently received much attention, in terms of yield (Knapp & van der Heijden,2018), of effects on biodiversity (Caprio et al., 2015; Lourenço et al., 2021; Smith et al., 2020), of nutritional value and of global positive impact on human health (Gomiero, 2018; Salomé et al., 2021).

Pest management in OF relies on specific cultural practices and a restricted number of non-synthetic pesticides. Conservation Biological Control (CBC) methods that enhance natural enemy abundance and activity to reduce pest populations (Heimpel & Mills, 2017; Holland et al., 2016) are of particular interest for OF. A wealth of literature, however, demonstrates that the potential and efficiency of CBC within a field do not only depend on local agricultural practices but also on the structure of the surrounding landscape (Begg et al., 2017; Muneret, Auriol, Thiéry, et al., 2019), including its amount of semi-natural habitat (Le Gal et al., 2020; Zamberletti et al., 2021) and the characteristics of the crop mosaic (Hillaert et al., 2018, 2020; Le Gal et al., 2020; Smith et al., 2020).

Organic farming has been expanding recently around the world in response to growing consumer demand and environmental concerns, and the area under OF is expected to increase in the future (Paull & Hennig, 2016). A majority of the literature on OF expansion points toward new organic farmers and fields clustering around existing ones (Gabriel et al., 2009; Marton & Storm, 2021; Sánchez Herrera & Dimitri, 2019; Zollet & Maharjan, 2021). Clustering happens for socio-economic and agronomic reasons, because OF conversion happens primarily “in agriculturally less-favored areas where economic incentives for conversion to organic farming do not need to be high and the loss of production due to conversion will be comparatively small” (Gabriel et al. 2009) such as isolated, hard to access, or less productive areas (Ilbery et al., 1999), but also because OF is often driven by newcomers, who could settle down more easily in the proximity of existing clusters, in less-favored areas, and in places where a “prevalence of small-scale, part-time, and self-sufficiency-oriented farming” is observed (Zollet & Maharjan, 2021).

The effect of OF expansion on pests and their natural enemies can be approximated, in a space for time approach (Blois et al 2013), by investigating how pests and natural enemies are affected by the proportion of OF in the landscape. Studies that investigated pest abundance in crops as a function of OF area in the landscape found that pests were either less (Gosme et al., 2012) or similarly (Muneret et al., 2018; Ricci et al., 2009) abundant when OF area increased. Moreover, predators of pests were either more or similarly abundant (Diekötter et al., 2010, 2016; Djoudi et al., 2018, 2019; Inclán et al., 2015; Lefebvre et al., 2016; Muneret, Auriol, Thiéry, et al., 2019; Puech et al., 2015), reviewed in (Petit et al., 2020), suggesting that earlier studies showing increases in crop damage associated with OF may have been influenced by the low amount of OF in the landscape in its early beginnings and that OF expansion scenarios may be of maximum importance (Muneret, Auriol, Bonnard, et al., 2019).

The effect of OF expansion on pest abundance and CBC in organic and conventional fields is difficult to predict. It will depend on the abilities of the pests and predators to develop in organic and conventional fields, on the interplay between pest and predator and landscape structure that conditions the ability of pests and predators to move among crops and semi-natural habitats (Kremen et al., 2007; Le Gal et al.,2020). More complex landscapes, i.e. landscapes with more, and more fragmented, semi-natural habitats and a more heterogeneous crop mosaic, may sustain more biodiversity (Batáry et al., 2011; Petit et al.,2020; Smith et al., 2020; Tscharntke et al., 2021; Tuck et al., 2014) and may support higher spill-over of predators from semi-natural habitats into crops (Concepción et al., 2008; Tscharntke et al., 2012). Such landscapes may also exhibit more movements of pests from semi-natural habitats to crops if pests find resources in semi-natural habitats at some point of their life cycle (Juhel et al., 2017). As a result, an increasing amount of semi-natural habitat in the landscape generally increases the abundance and diversity of natural enemies as well as pest predation and parasitism but its effect on pest abundance or damage is case dependent (Chaplin-Kramer et al., 2011; Karp et al., 2018; Veres et al., 2013). Similarly, pest and predator movements between organic and conventional crops are expected to increase with the edge length between these two crop types. Organic expansion should thus affect more pest abundance in conventional crops when the two crop types are interspersed. The response of pest abundance to OF expansion may moreover differ in organic and conventional fields: local management is expected to have large effects on biodiversity or ecological functions when landscapes are of intermediate intensity but to have little effect when landscapes are either very or very little intensive (the intermediate landscape hypothesis (Perez-Alvarez et al., 2019; Tscharntke et al., 2005, 2012)) and, reciprocally, landscape effects are supposed to depend on local practice intensity (Petit et al., 2020). Such interactions have, however, seldom been observed in the field (Petit et al. 2020, but see e.g. Perez-Alvarez, Nault, et Poveda 2019;Ricci et al. 2019).

Given the inherent complexity of conservation biological control (Begg et al., 2017) and the lack of CBC data in the context of OF expansion, modeling appears as a useful tool to understand and forecast how pests and their control may vary during OF expansion in a diversity of landscape contexts. The only published modeling study to our knowledge considered a pest-parasitoid system in a landscape exclusively composed of conventional and organic fields (Bianchi et al., 2013). This study interestingly showed that clustering organic and conventional fields decreased the proportion of OF necessary for maintaining the parasitoid population and decreased pest load. It also showed that intermediate levels of OF may lead to transitory peaks in pest load due to the delay of the parasitoid population response to pest abundance (Bianchi et al., 2013). It is therefore interesting and necessary to study, through modeling approaches, how spatial scenarios of organic farming expansion impact conservation biological control (Adl et al., 2011; Bianchi et al., 2013). As stated by Petit et al, (2020), such modeling approaches “can offer in silico tests of the consequences of much larger proportions of agroecological practices in the landscape” and could be combined with empirical studies to “provide key insights about how natural enemies and pests will behave in future landscapes.”

In the following, we pair a grid-based landscape model and a spatially explicit Lotka-Volterra type predator-prey model (Ciss et al., 2016; Roques, 2015) to investigate how contrasted scenarios of OF expansion, defined by their spatial arrangements, impact pest abundance in organic and conventional crops. The scenarios are applied to a diversity of landscapes differing in their amount of semi-natural habitat and its fragmentation. We further investigate if the ranking of scenarios is robust to pest control methods in OF fields and pest and predator dispersal abilities.

## Material and Methods

### 1. Overview

The modeling procedure comprises three main elements. The first is a stochastic landscape model to initiate the structure of the landscape, i.e. the total area and fragmentation of semi-natural habitat and the initial area of organic and conventional crops. The second is a population dynamics model to represent the dynamics of interacting pests and their predators on the changing landscapes. The third is a set of spatial scenarios of OF expansion that govern landscape change over time (Figure 1).

**Figure 1.**
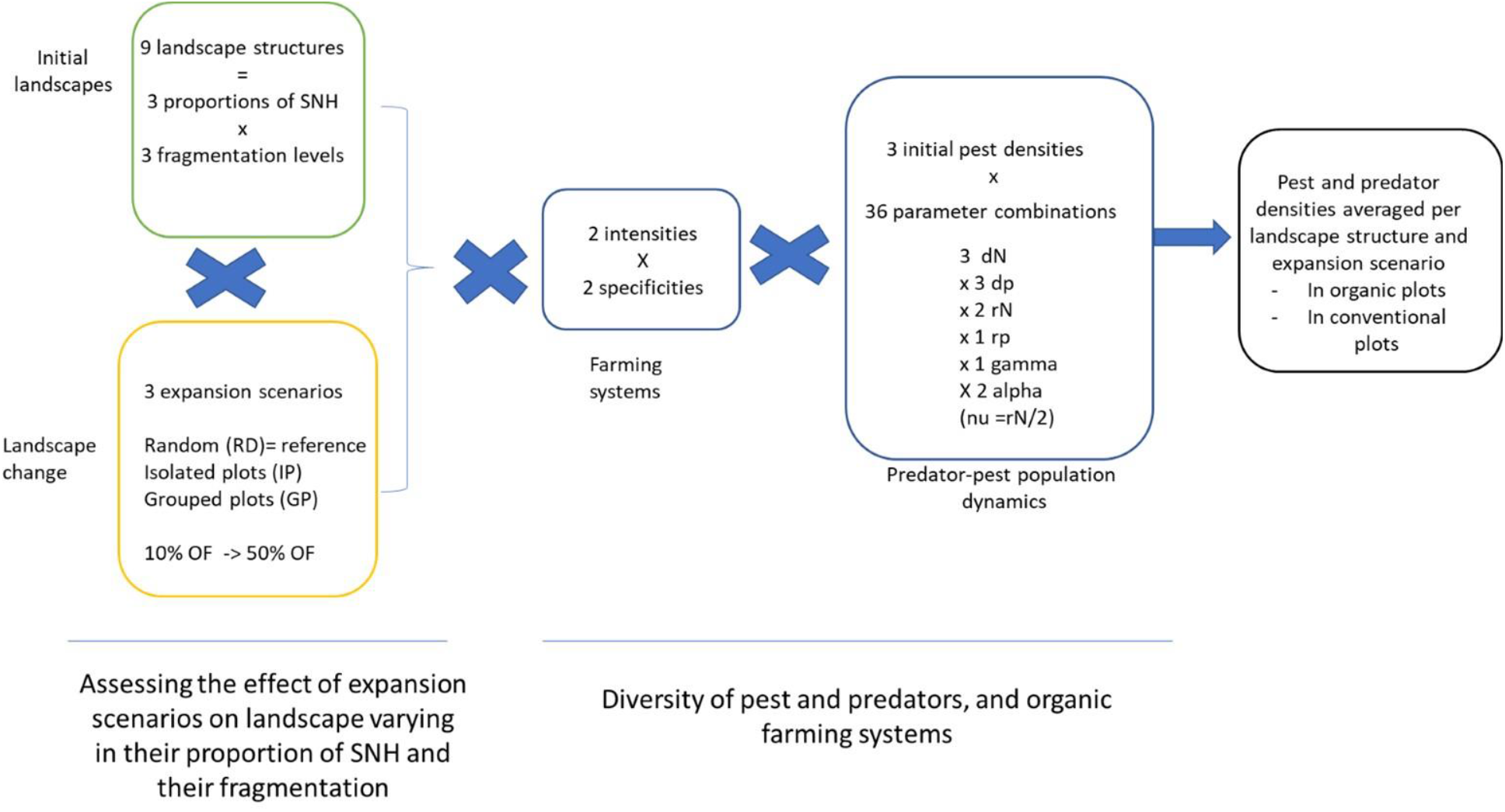
The simulation design combines three spatial scenarios of organic farming expansion (Random versus IP or GP) in nine landscape contexts (3 proportions of seminatural habitats (SNH) x 3 fragmentation levels) for various predator-pest population dynamics (36 pest biology parameter combinations and 4 pest management types in the organic farming system). The green box corresponds to the landscape model, the blue box to the population dynamic model and the orange box to the land change scenarios.

**Figure 2.**
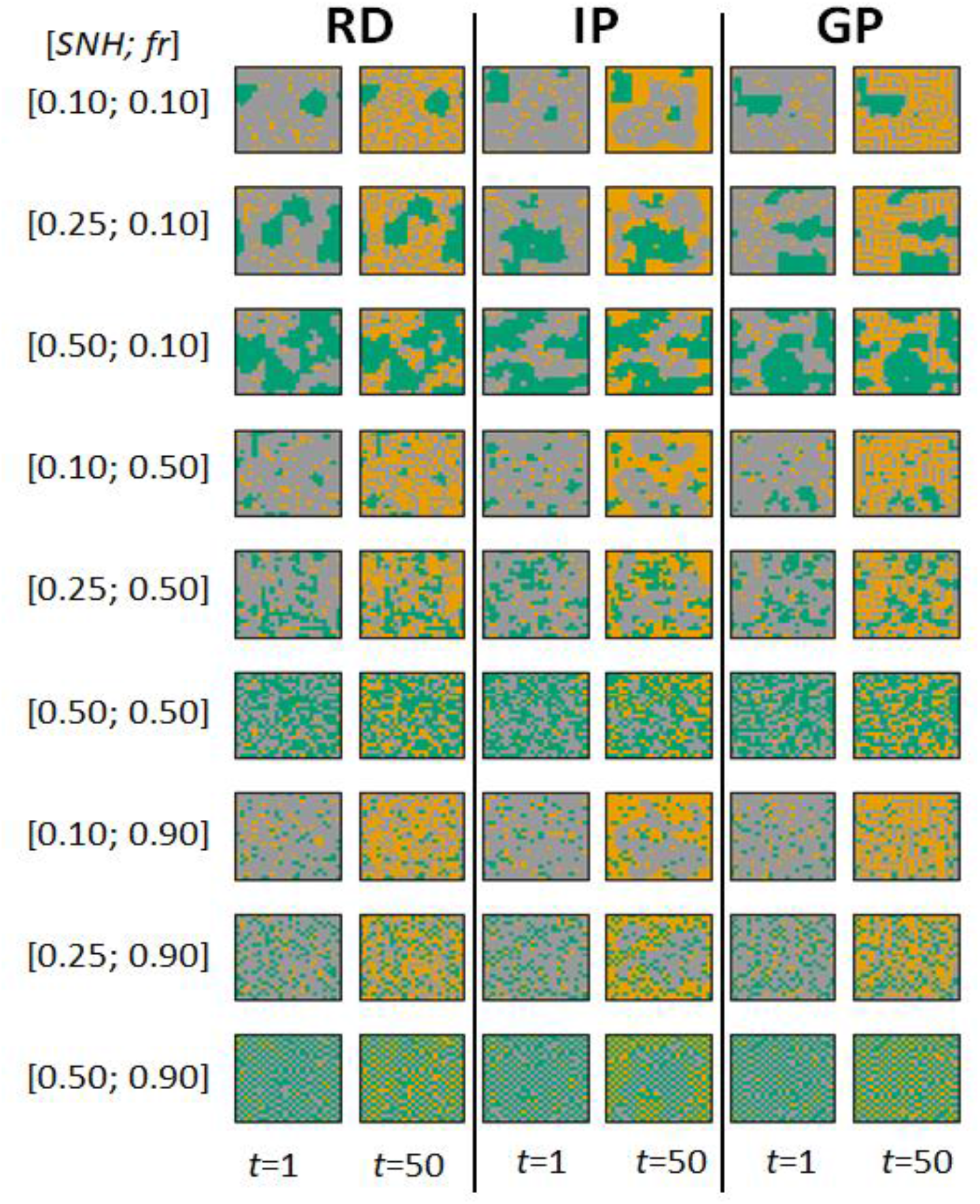
Examples of landscape structures at the beginning (t=1) and the end (t=50) of the organic expansion. The figure provides one example landscape for each combination of landscape context and OF expansion scenarios. SNH= proportion of semi-natural habitat; *fr:* fragmentation of semi-natural habitat; RD, IP and GP refer to the three scenarios of selection of CF fields to convert to OF: selection of random, isolated or grouped fields; Grey: conventional (CF), orange: organic (OF), green: semi-natural habitat (SNH).

### 2. Landscape and population dynamics models

#### 2.1. Stochastic landscape model to set the initial landscape context

To set the landscape context in which to initiate the joint simulation of OF expansion and pest and predator population dynamics, we used a landscape model initially developed by Roques and Stoica (2007), then improved by Roques (2015). This landscape model generates stochastic landscapes with several types of land uses over a square grid (*n × n* matrix). Land use allocations were parametrized by the proportion of each type of land use and the fragmentation level of one target land use (Roques 2015;Ciss et al. 2016). This fragmentation is controlled by the *fr* parameter, which is an index of fragmentation *per se* (Fahrig, 2003). *fr* measures the landscape-level average proportion of neighbors among the 4 closest neighbors of each cell of that land use that are not of the same land use type. *fr* is in the range [0,1], and increases with fragmentation. To reach the desired fragmentation level, grid cells are first randomly allocated to one of the land uses, controlling for the proportion of cells associated to each land use. Then a Metropolis-Hastings algorithm is run to optimize the spatial distribution of the cells associated with the target land use. The algorithm stops when the distance between the observed and the desired fragmentation index is below a tolerance threshold.

Here, we set up initial landscapes composed of three land uses: (i) semi-natural habitats (SNH), (ii) Organic Farming crops (OF), and (iii) Conventional Farming crops (CF). We controlled the proportions of these land uses and the fragmentation level *fr* of SNH. (Table 1). *fr* is thus an index of fragmentation *per se*. High values of *fr* resulted in landscapes with more numerous and smaller SNH patches and increased edge length between SNH and crop patches (Fig. 1, S1.1, S1.2, S1.3).

**Table 1.** List of model parameters.

| Parameter | Description | Unit | Values |
| --- | --- | --- | --- |
| **Landscape model:** |  |  |  |
| $n$ | Size of the lattice | | 24 |
| $fr$ | Fragmentation index | dimensionless | {0.1,0.5,0.9} |
| $pSNH$ | Percentage semi-natural habitat | % | {10, 25, 50} |
| T | Time horizon | year | 50 |
| **Model of population dynamics:** |  |  |  |
| $d_N$ | Pest dispersal coefficient | unit area.year<sup>-1</sup> | $\frac{1}{n^2}\{0.1,0.5,1\}$ |
| $d_P$ | Predator dispersal coefficient | unit area.year<sup>-1</sup> | $\frac{1}{n^2}\{0.1,0.5,1\}$ |
| $r_N$ | Pest intrinsic growth rate | year<sup>-1</sup> | $2\{\ln(50), \ln(100)\}$ |
| $r_P$ | Predator intrinsic growth rate | year<sup>-1</sup> | $\ln(2)$ |
| $\gamma$ | Predator life expectancy in crops | year | $1/2$ |
| $\alpha$ | Predation index | indiv<sup>-1</sup> year<sup>-1</sup> | $\frac{1}{\gamma}\{4/3, 4\}$ |
| $v$ | Pest management effect | year<sup>-1</sup> | $r_N/2=\{\ln(50), \ln(100)\}$ |

#### 2.2. Population dynamic model

##### 2.2.1 General description

We modeled the spatio-temporal dynamics of a pest and a generalist predator species interacting over the lattice generated by the landscape model according to (Martinet & Roques, 2022). The model describes the density of the predator population *P_t_* (*x*) and of the pest population *N_t_* (*x*) at each position *x* = (*i, j*) over the grid and at each time step *t* (equations 1). The variation over time (indicated with sign ‘) of pest (*N′_t_* (*x*)) and predator (*P′_t_* (*x*)) densities at each position depends on their dispersal in and out of this position, their intrinsic growth (i.e. population growth in absence of pesticides and of interactions between pests and predators), mortality due to pesticides, and mortality (for the pest) or growth (for the predator) due to predation.

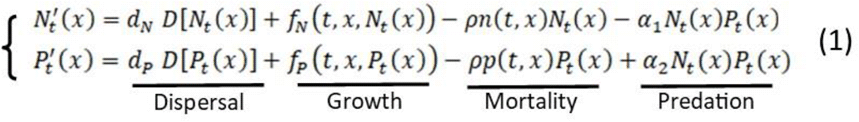

###### Dispersal

*D*[.] defined as 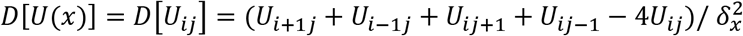 is the discrete Laplace operator modeling the movements of the individuals to adjacent cells, *δ_x_* = 1/*n* being the length of a unit cell in the landscape. From each position *x* = (*i,j*), and during a time interval *δ_t_* ≪ 1, a proportion 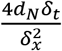 of the pest population (resp. 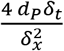 of the predator population) moves to its 4 adjacent cells. Thus *d_N_* and *d_P_* directly control the mobility of the pest and predator populations. We assume periodic conditions at the boundaries of the lattice.

###### Growth

The terms *f_N_*(*t*, *x*, *N_t_* (*x*)) and *f_p_*(*t*, *x*, *P_t_* (*x*)) stand for the pest and predator intrinsic growth functions. They are controlled by parameters *r_N_* and *r_p_* respectively (Table 1). The predator being a generalist, it can grow in absence of pests.

###### Mortality

*ρ_N_*(*t, x*)*N_t_* (*x*) and *ρ_P_*(*t, x*)*P_t_* (*x*) account for the pest and predator death rates caused by pest management. Mortality is controlled by parameter *ν* (Table 1).

###### Predation

The interaction terms –*α*_1_*N_t_*(*x*)*P_t_*(*x*) and *α*_2_*N*(*x*)*P_t_*(*x*) describe the effects of predation on the pest and predator growth rates, respectively. We assume standard Lotka–Volterra interactions between the pest population and its predator, which means that the pest death rate increases linearly with the density of the predator, and conversely the growth rate of the predator increases linearly with the pest population density. We also assume that *α*_1_ = *α*_2_ = *α* (Table 1)

The system is scaled so that the carrying capacities of *P* and *N* are both equal to 1 thus the population densities are expressed in units of their respective carrying capacities.

##### 2.2.2 Timing of ecological processes

The year is divided into equal intervals *δ_t_* each corresponding to a dispersal event of pests and predators. The number of intra-annual time steps is thus calculated as 1+ (1/*δ_t_*). Each year is divided into two periods [0, ½[and [½, 1] during which ecological processes differ (Table 2). Pests are specialized on the crop and their phenology matches that of the crop. The first half of the year schematically represents the season where the crop is absent, pests do not reproduce and there are no pest management practices. Pest densities only depend on their dispersal and predation by predators. During the second half of the year, when the crop is present, pests furthermore reproduce and are affected by pest management practices in the crop. The predators, in contrast, reproduce all year long in semi-natural habitats (loosely mimicking taxa that include both spring and autumn reproduction) and suffer from intrinsic mortality in crops. Their density increases further in both semi-natural habitats and crops when the pest is present. Predators thus behave as generalist predators that feed on the pest prey, and on alternative prey in semi-natural habitats. Like pests, they are affected by pest management practices in crops during the second half of the year.

**Table 2.** Values of the growth functions *f_N_*(*t, x, N_t_*(*x*)) and *f_P_*(*t,x, P_t_*(*x*)). *r_N_* is the pest intrinsic growth rate in the crops in the absence of pest management, *γ* the natural enemy life expectancy in the absence of resources and *r_P_* the natural enemy birth rate in semi-natural habitats.

| Time span | $f_N(t, x, N)$ | | $f_P(t, x, P)$ | |
| --- | --- | --- | --- | --- |
| | $[0, \frac{1}{2}[$ | $[\frac{1}{2}, 1]$ | $[0, \frac{1}{2}[$ | $[\frac{1}{2}, 1]$ |
| **Land use** |  |  |  |  |
| Conventional | 0 | $r_N N(1 - N)$ | $-P/\gamma$ | $-P/\gamma$ |
| Organic | 0 | $r_N N(1 - N)$ | $-P/\gamma$ | $-P/\gamma$ |
| Semi-natural habitats | 0 | 0 | $r_P P(1 - P)$ | $r_P P(1 - P)$ |

##### 2.2.3 Organic farming systems

There exists a diversity of organic farming systems with more or less intensive pest management strategies (Marliac et al., 2015)). To represent this diversity, we considered four theoretical types of organic farming (Table 3). In the most intensive OF systems *(Int-Gen* and *Int-Spe*), pest management is as efficient in organic fields as in conventional ones so that the mortality of pests due to pest management practices is the same. These two OF systems differ by the specificity of these control measures that either do not *(Int-Spe)* or do *(Int-Gen)* affect predators, but less than in conventional fields. Examples of efficient and specific pesticides are microorganisms targeting pests such as viruses (Graillot et al., 2016) or other microorganisms (Deshayes et al., 2017). Other pest management measures such as pesticides *(e.g.* spinosad) or nets are efficient on pests but also affect some predators (Dib et al., 2010).In the extensive OF systems *(Ext-Gen* and *Ext-Spe)* pest management is less efficient in OF fields and pest mortality rates are half those in conventional fields. As before, these two OF systems differ by the specificity of their pest management practices that either do *(Ext-Gen)* or do not *(Ext-Spe)* affect predators (Table 3).

**Table 3.** Effects of pest management practices on the pest and the natural enemies in conventional (CF) and organic farming. Four organic farming systems were considered. Int-Spe and Int-Gen correspond to intensive pest management (high pest mortality) while Ext-Spe and Ext-Gen are less intensive. In Int-Spe and Ext-Spe systems, pest management practices are specific to the pest and do not affect predators while in Int-Gen and Ext-Gen systems, management is less specific.

| | $\rho_N(t, x)$ | $\rho_P(t, x)$ |
| --- | --- | --- |
| **CF** | $2\nu$ | $2\nu$ |
| **Int-Spe** | $2\nu$ | 0 |
| **Int-Gen** | $2\nu$ | $\nu$ |
| **Ext-Gen** | $\nu$ | $\nu/2$ |
| **Ext-Spe** | $\nu$ | 0 |

##### 2.2.4 Parameter values

###### Intrinsic growth rates

The pest reproduces only during the second half of the year. During one year, the population would increase by a factor of exp(rN)/2 in the absence of any limiting factor. We thus assumed that, under these conditions, the population would increase by a factor of 50 or 100 over the season. We assumed a lesser yearly increase for the predator of exp(rp)=2, i.e. a population doubling in the absence of pests or any limiting factor. To compensate for this increase, we assumed a life expectancy of the predator on the crops of *γ*=1/2 year, in the absence of pests.

###### Mortality due to pest management practices

We assumed that the mortality rate induced by pest management practices is comparable to pest growth rates (2 *ν*∈2{ln 50, ln 100 }). The mortality due to pest management practices reaches its maximum value for both pests and predators in the conventional fields and for pests only in the most intensive OF systems (Int-Spe and Int-Gen). In these situations, mortality compensates for the pest population’s local increase and drastically reduces predator populations. Mortality caused by pest management practices is reduced by half or set to 0 for predators depending on the OF systems (Table 3).

###### Dispersal

The values for *d_N_* and *d_P_* were chosen so that approximately between 0.1% (*d_N_* or *d_P_*=0.1/n^2^) and 1% (*d_N_* or *d_P_*= 1/n^2^) of individuals in a given cell move to neighboring cells every day.

### 3. Initial Conditions

#### 3.1. Landscapes

Simulations were run on 9 landscape contexts differing in their proportion of semi-natural habitats (SNH) (either 10, 25, or 50% of total area) and in the fragmentation of these habitats (*fr* values: 0.1, 0.5 and 0.9). Initially, 10% of crops were organic (OF) and 90% conventional (CF) based on the current national proportions in France (ORAB PACA, 2020) and on the proportion of worldwide cropped and pasture land that is practicing some forms of organic farming (Pretty et al, 2018). Based on that, we generated initial landscapes with three proportions of each land-use, named respectively Qin1 (10% SNH; 9% OF; 81% CF), Qin2 (25% SNH; 7.5% OF; 67.5% CF), and Qin3 (50% SNH; 5% OF; 45% CF). In the remainder of this paper, we refer to these three initial conditions in terms of their SNH proportions (SNH 10%, 25% and 50%). Initial OF crops were allocated randomly among crop cells. Each simulation of the model was run on a different initial landscape.

#### 3.2. Population dynamics

At t=1 predators are introduced in all semi-natural habitats with initial density P1_SNH_. The predators are allowed to reproduce and disperse until t=3. At t=3 pests are introduced in the crops with initial density N1_crop_ = P1_SNH_. To assess the impact of initial conditions on our conclusions, we set three extreme values for P1_SNH_: 0.1, 0.5 and 1. We then performed simulations during a 15-years burn-in period in order to allow the stabilization of pest and predator dynamics before organic farming expansion.

### 4. Spatial scenarios of organic farming expansion

From each initial landscape, we simulated OF expansion from t=15 to t= 50 years in order to sequentially convert 50% of the initial conventional crop area to OF. For each simulation, OF expansion was progressive, i.e. approximately 6.25% of the initial conventional crop area was converted to OF every 5 years. The total number of conventional fields to be converted depended on the initial cultivated area and the target proportion of OF. The final compositions of landscapes corresponding to the three initial proportion of semi-natural habitats were respectively Qen1(10% SNH; 49.5% OF; 40.5% CF), Qen2(25% SNH; 41.25% OF; 33.75% CF), Qen3(50% SNH; 27.5.5% OF; 22.5% CF). Only conventional fields were converted to organic. The area of semi-natural habitat remained constant.

Three spatial conversion scenarios were simulated based on the 4-neighborhood of conventional fields:

- the RD scenario in which we performed a random choice of conventional fields to be converted,
- the IP scenario, in which isolated conventional fields, i.e. fields with as few as possible conventional 4-neighbors, were converted first,
- the GP scenario in which, in contrast to IP, conventional fields with as many as possible conventional 4-neighbors were converted first.

The IP and GP scenarios are two possibly planned scenarios that we compared to the baseline RD scenario in terms of resulting pest densities and predator to pest ratio.

### 5. Simulation outputs

At each time step of each simulation, we recorded indicators of the landscape structure and of pest and predator densities in each land use (CF, OF and SNH).

#### 5.1. Landscape structure

Landscapes can be described in terms of composition, i.e. proportion of the land uses, and configuration, i.e. the spatial arrangement of these land uses (Fahrig & Paloheimo, 1988). Landscape composition was controlled during the simulation. We monitored landscape configuration using three landscape metrics for each land use: the mean patch area, the number of patches, and the edge length (R package *landscape metrics,* Hesselbarth et al, 2019). For a given land use, patches were made of fields of that given land use that were 4-neighbors to at least one field of the same land use. Together, these three metrics indicate whether, for a given proportion of landscape area, one land use is represented by a few large patches or many small patches.

#### 5.2. Pest and predator densities

For each simulation, the densities of pests and predators were monitored at the end of each year and averaged over each land use (SNH, OF and CF). From these, a median predator to pest ratio was calculated per land use as a proxy of the intensity of pest control by predators.

### 6. Simulation study

Simulations for the three spatial organic farming expansion scenarios mentioned above were performed for each of the nine types of landscapes (3 proportions of SNH x 3 levels of *fr)* aiming at 50% OF fields for each of the four types of OF (Table 3). These simulations were performed for all combinations of the values of the 6 parameters (pest and predator dispersal coefficients, pest and predator intrinsic growth rates, predator life expectancy in crops, interaction term,) and the four farming systems governing pest and predator population dynamics (Table 1, Fig. 1) and the three initial densities of pests and predators. This resulted in a total of 11664 Simulations, each run on a different landscape. We performed 11664 more simulations without any action on the landscapes. These simulations are referred to as Reference (REF).

Comparisons of pest and predator densities and predator to pest ratios among conversion scenarios were performed at the end of the simulations (t=50) for each landscape context. As pest density was the main variable of concern regarding OF expansion, we further checked whether the ranking of scenarios was robust with regards to the intensity of OF and the dispersal rate of the pest.

All simulations were performed with (*MATLAB,* 2018a) and all statistical analyses were performed with R software (*R Software,* 2017).

## Results

### 1. Pest and predator dynamics

#### 1.1 Pest and predator dynamics in absence of organic farming expansion

The model behavior was first studied in the absence of organic farming expansion, i.e. at the initial proportions of organic farming. This step first shows that in the absence of organic farming expansion, the average landscape scale densities of pest and predator remained stable over time after approximately 15 years (Figure 3, scenario: REF). These equilibrium densities were independent of the initial pest density (not shown). Consistent with parameter values, both pest and predator densities were higher in organic fields than conventional fields. Moreover, the density of pests was always larger than that of predators, in both conventional and organic fields (Fig. 3, scenario: REF).

**Figure 3.**
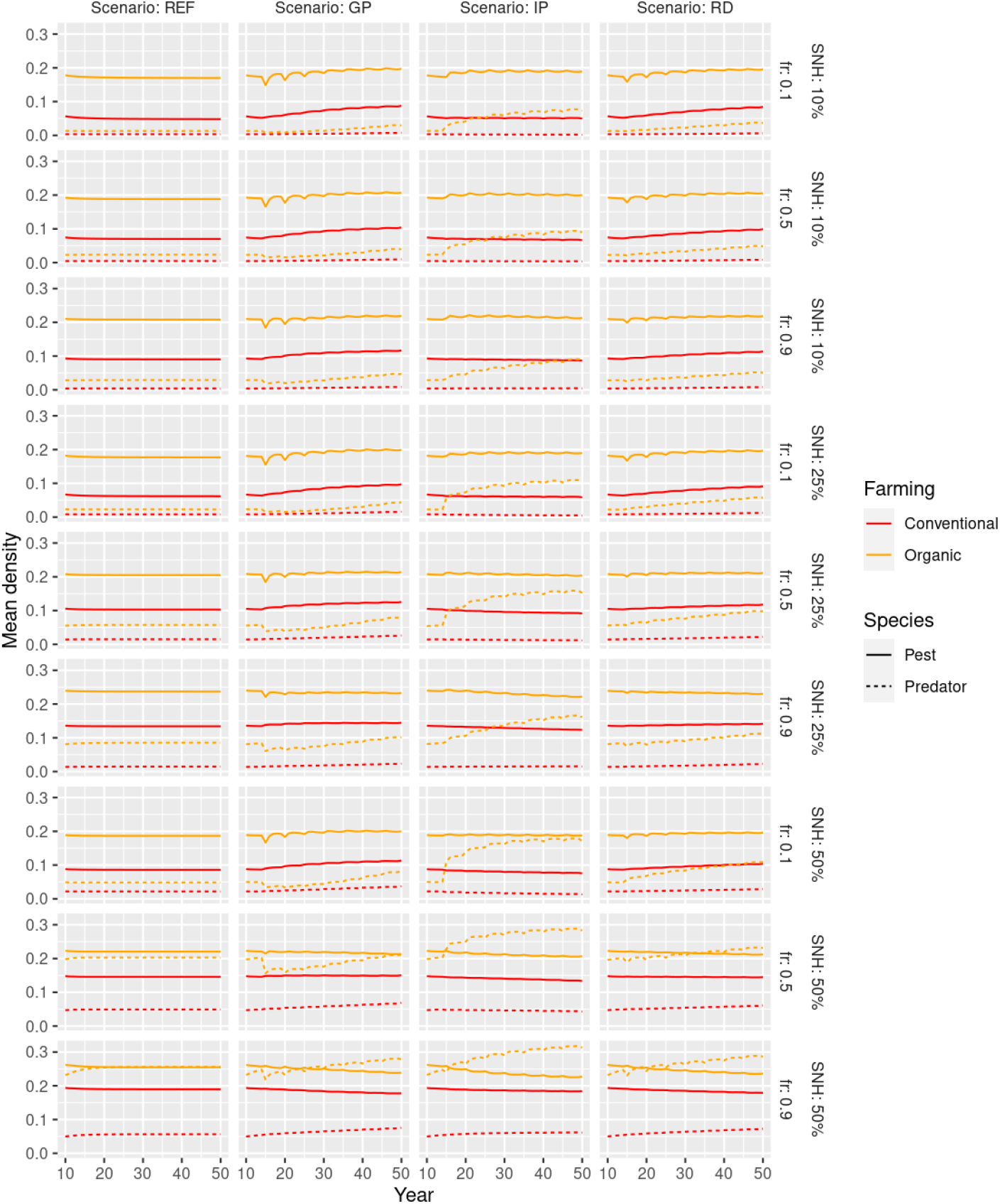
Effect of organic expansion scenarios on the temporal evolution of the mean density of pests and predators in organic (OF) and conventional farming (CF), as a function of fragmentation *(fr)* and initial proportion of semi-natural habitat (SNH). Reference (REF) represents the absence of organic expansion.

Pest and predator densities did not show a clear response to the increase of proportion of semi-natural habitat and they increased with its fragmentation, probably because cultivated fields were more likely to be close to a semi-natural habitat, increasing spill-over of individuals into cultivated fields. These effects were stronger on predators than pests in organic fields, consistent with the higher dependency of predator reproduction and survival on semi-natural habitats. The predator density, in contrast, remained very low in conventional fields due to pesticides.

These differential effects of landscape characteristics on pests and predators had two consequences. First, pest densities were 2.6 times larger in organic fields than in conventional fields in landscapes with little semi-natural habitat and little fragmentation (SNH=10%, fr=0.1) while they were only 1.3 times larger in landscapes with large proportions of fragmented semi-natural habitats (SNH=50%, fr=0.9). Second, the predator to pest ratio increased in organic fields but decreased in conventional fields when semi-natural habitat proportion and fragmentation increased.

#### 1.2 Pest and predator dynamics during organic farming expansion

The cultivated landscape changed during organic farming expansion. Compared to their initial area, at the end of organic farming expansion, conventional patches were generally smaller and organic patches larger. Constraints were furthermore imposed by the spatial distribution of semi-natural areas so that patch area varied more in little fragmented landscapes or when there was little semi-natural area. Because they set different priorities regarding field conversion, the different scenarios led to different cultivated patch area dynamics, with some dramatic changes due to conventional patch splitting. The IP scenario notably always resulted in conventional patches that were larger than the other scenarios while the GP scenario generally resulted in larger organic patches (Supplementary material S1.3).

The organic farming expansion affected more predator densities than pest densities for most combinations of landscape contexts and expansion scenarios (Fig. 3). Its impact was also generally stronger in organic than conventional fields and in landscapes with large proportions of fragmented semi-natural habitats. As expected, predator densities in organic fields generally increased. Changes in pest and predator densities and their dynamics, however, depended on expansion scenarios and landscape contexts.

##### 1.2.1 Pest dynamics

Pest densities in organic fields showed similar changes for the three expansion scenarios (Fig. 3). They tended to slightly increase or remain stable over time in landscapes with little or intermediate proportions of semi-natural habitat and to decrease in landscapes with large proportions of semi-natural habitat. In conventional fields, pest densities showed this same pattern with the RD and GP scenarios but not with the IP scenario. With the IP scenario, pest densities in conventional fields tended to decrease slightly over time whatever the landscape context. As a result, at t=50, pest densities were generally smaller with the IP than with the RD and the GP scenario in conventional fields and similar for the three expansion scenarios in organic fields.

##### 1.2.2 Predator dynamics

In organic fields, the effect of organic farming expansion on predator densities was very large compared to its effect on pest densities (Fig. 3). Predator densities increased for the three expansion scenarios. The increase was larger for the IP scenario than for other scenarios, particularly in little fragmented landscapes with intermediate or large proportion of semi-natural habitats. For example, when SNH=25% and *fr*=0.1, with the IP scenario the predator density at t=50 was 5.38 times larger than the initial density and was 2.44 times higher than the predator density at t=50 with the GP scenario. In contrast, the three scenarios performed similarly in landscapes with the highest proportion and fragmentation of semi-natural habitat (SNH=50%, fr=0.9). In these landscapes, the predator density increased by a factor of 1.34 between t=0 and t=50 with the IP scenario and was only 1.20 times higher than with the GP scenario at t=50. The increase in predator density was moderate for the RD and GP scenarios and reached similar values at t=50. Their dynamics were, however, qualitatively different. While predator densities increased steadily for the RD scenario, for the GP scenario, most predator densities showed a transient decrease in the first years following the beginning of organic farming expansion.

Note that in landscapes with 50% SNH predator densities were sometimes larger than pest densities in organic fields (Fig. 3). This was most prominent when fragmentation was high, an indication that it resulted from spillover of predators from semi-natural habitats.

The pattern was very different in conventional fields. Predator densities remained stable at very low values for most landscapes and expansion scenarios. They only increased in the GP scenario in landscapes with high proportion of semi-natural habitats but still remained at low values.

### 2 Effect of spatial scenarios of organic farming expansion and landscape contexts on resulting pest densities and conservation biological control

#### 2.1 Pest densities

In organic fields, differences in final pest density were limited among expansion scenarios. Pest density in organic fields responded overall little to landscape characteristics and, in particular, less to the different scenarios of OF expansion, despite differences in organic or conventional patch areas (Fig. S1.3), than to the fragmentation of semi-natural habitats (Fig. 4, upper panel). The highest levels of pest densities were obtained for the highest fragmentation levels. For a given level of fragmentation, pest densities in organic fields tended to be lower for the IP scenario but the amplitude of effect was smaller than for fragmentation. In contrast, final pest density in conventional fields (Fig. 4, lower panel) responded both to the OF expansion scenario and to fragmentation, indicating a dependence on conventional and organic patch area (Fig S1.3). As in organic fields, pest density increased with the level of semi-natural habitat fragmentation. In conventional fields, low levels of pest densities could thus be attained for different fragmentation levels given that conventional patch areas were large, a situation provided by the IP scenario in landscapes with small proportion of semi-natural habitats (SNH=10%). Furthermore, the range of variation of pest densities was larger in conventional than organic fields.

**Figure 4.**
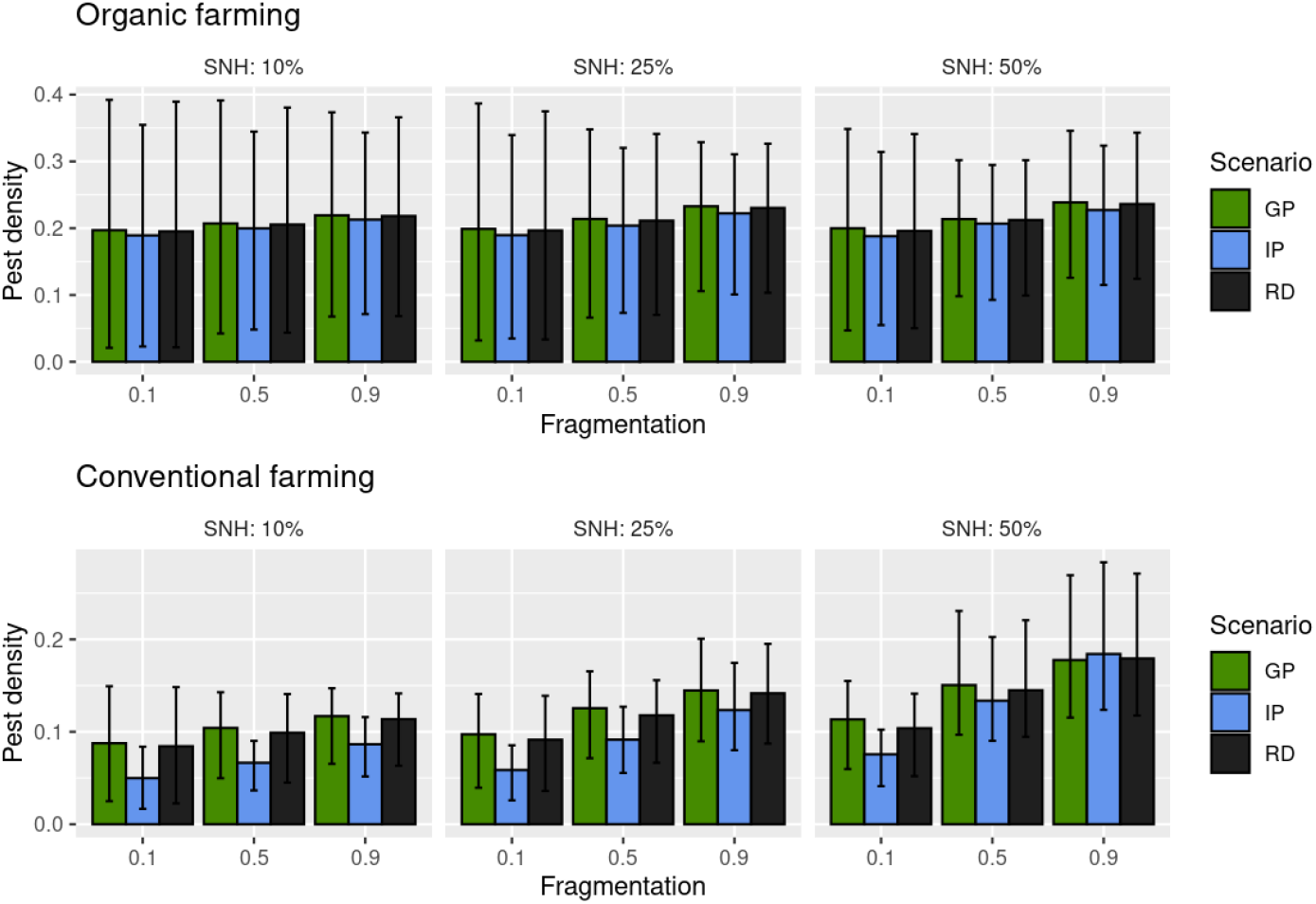
Mean final (t=50) density of pests in organic (upper panel) and conventional (lower panel) fields as a function of the landscape context (fragmentation and percentage of semi-natural habitat). Results are presented for each OF spatial expansion scenario: GP (grouped plots), IP (isolated plots), RD (random).

#### 2.2 Conservation biological control

The predator to pest ratio is an indicator of the potential for conservation biological control: a higher ratio indicates that pests are more likely to come across a predator. As a result of the pest and predator dynamics described above, the predator to pest ratio at the end of the simulation was three to four times larger in organic fields than in conventional fields (Fig. 5). It increased with the proportion of semi-natural habitat, in similar relative proportions in organic and conventional fields, from an average of approx 0.2 to 1.25 in organic fields and 0.05 to 0.35 in conventional fields, when the proportion of SNH increased from 10% to 50%. It also increased, but to a much lesser extent with SNH fragmentation. The only significant increase with fragmentation was for landscapes with large proportion of SNH (Fig. 5).

**Figure 5.**
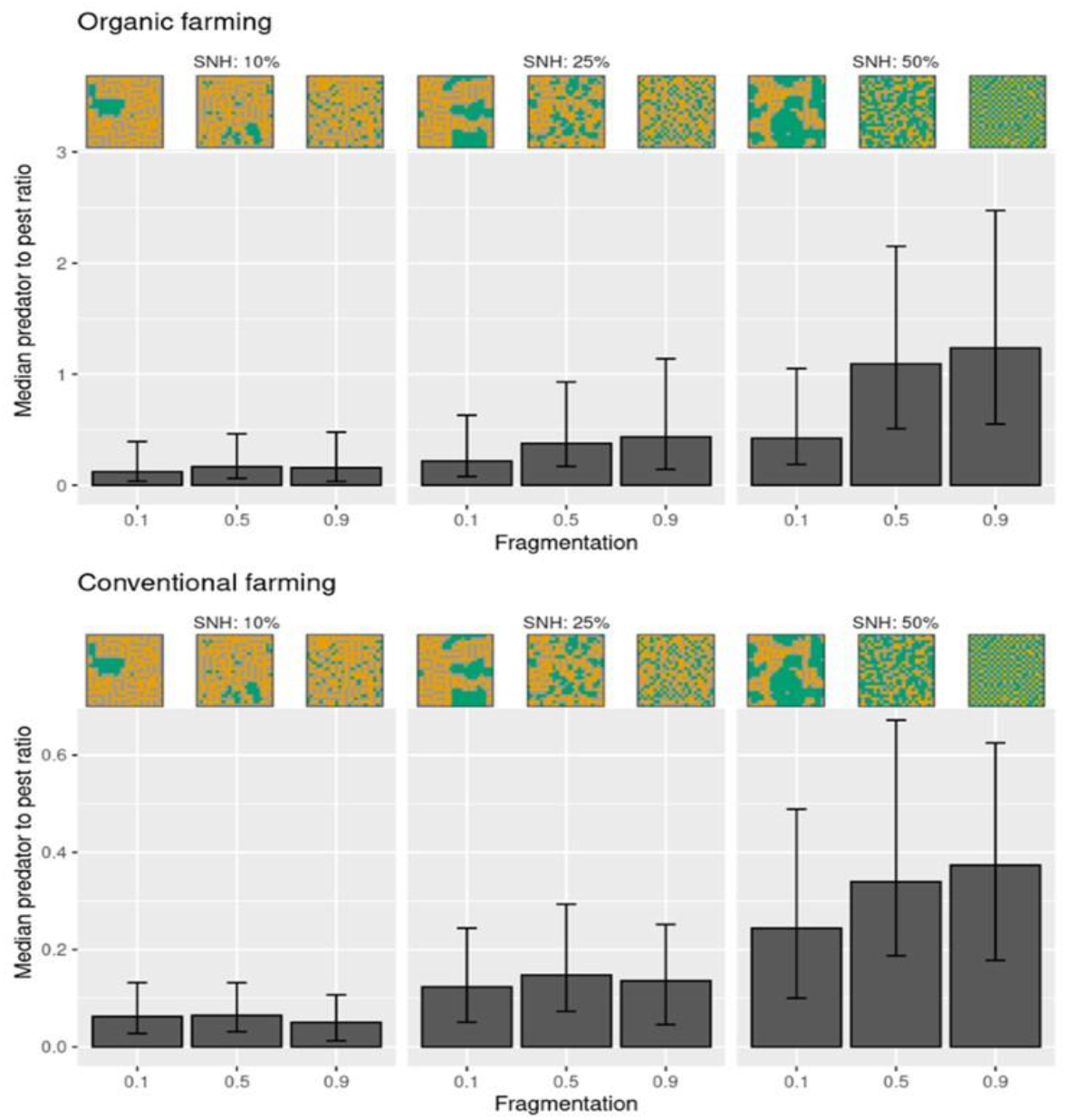
Median Predator to pest ratio at t=50 for the nine types of initial landscapes, in organic fields (upper) and conventional fields (lower). Error bars represent the first and third quartiles over all scenarios and parameter values. Examples of landscape contexts (combinations of fragmentation and percentage of SNH) are provided for illustration. Note that y-axes are on different scales.

More interestingly, we observed a clear ranking of spatial expansion scenarios with IP>RD>GP for the predator to pest ratio in organic fields (Fig. 6). This ranking might be due to the larger increase of predator densities during OF expansion with the IP scenario and the somewhat larger pest densities with the GP scenario (Fig. 3). Relatively to the RD scenario, the predator to pest ratio was from 1.83 times higher (SNH=10%, fr=0.1) to 1.1 (SNH=50%, fr=0.9) times higher for the IP scenario. In contrast, these ratios for the GP scenario ranged from 0.55 (SNH=10%, fr=0.1) to ^~^1(SNH=50%, fr=0.9) times those for the RD scenario.

**Figure 6.**
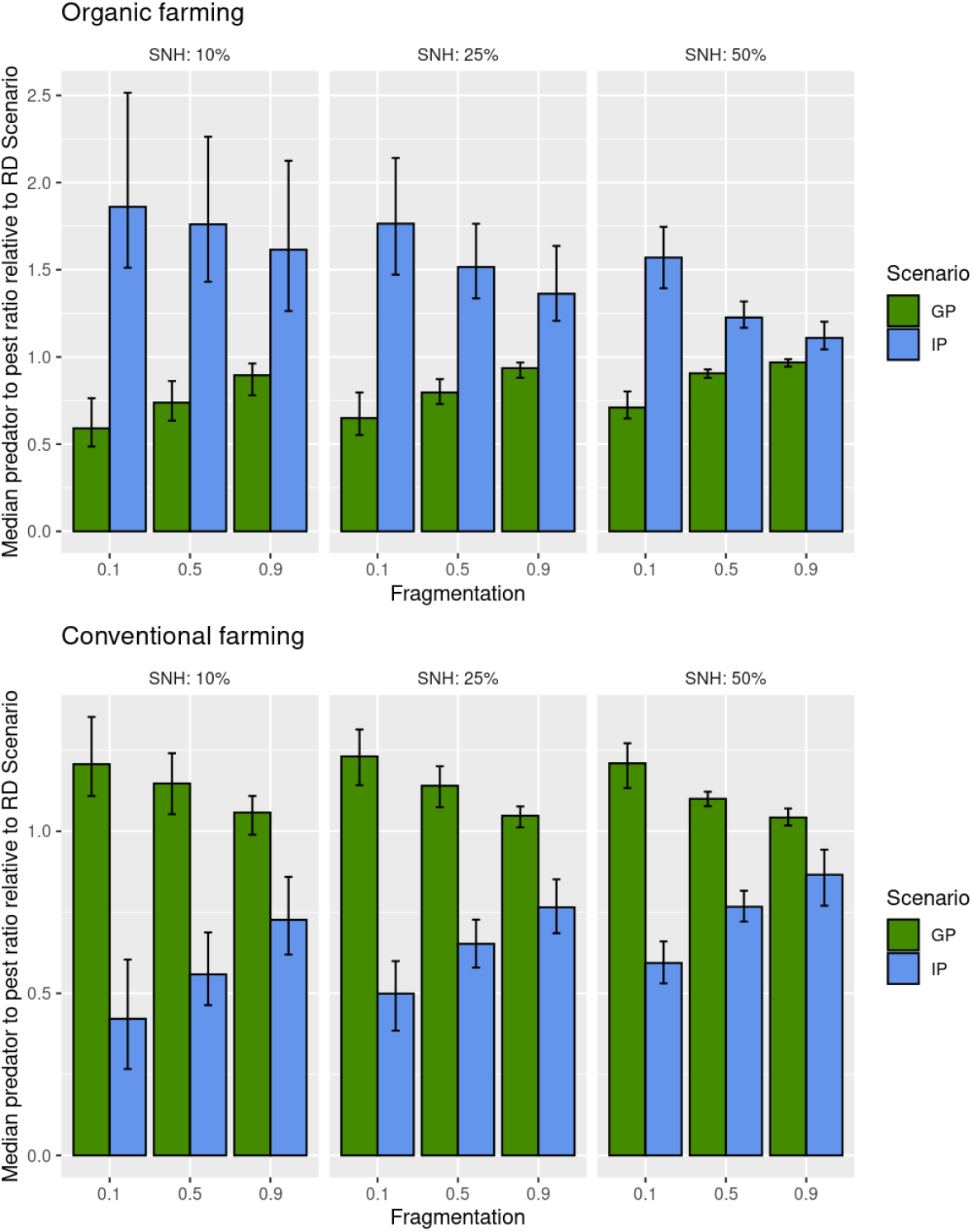
Median predator to pest ratio at t=50 in organic and conventional crops for the IP and GP scenarios relative to the ratio obtained for the RD scenario with the same parameter values. Error bars represent first and third quartiles over all combinations of population dynamic parameters.

In conventional fields, predator to pest ratios showed the opposite GP>RD>IP ranking. The difference here was mainly between the IP and the two other scenarios. Ratios were a little larger for the GP scenario than for the RD scenario whatever the landscape context, with values ranging from 1.2 (SNH=10%, fr=0.1) to ^~^1 (SNH=50%, fr=0.9) times those for the RD scenario. They were the smallest for the IP scenario, particularly in fragmented landscapes with low proportions of semi-natural habitats (from 0.55 (SNH=10%, fr=0.1) to 0.8 (SNH=50%, fr=0.9) times higher than with the RD scenario, Fig. 6). Consistent with the larger differences in crop patch area in landscapes with few and little fragmented semi-natural habitat, differences between the GP and IP scenarios were largest in such landscapes and very small in landscapes with a large proportion of highly fragmented semi-natural habitats.

## Discussion

Existing evidence of the positive impacts of OF (Organic Farming) on agrobiodiversity and pest control (eg. Muneret et al. 2019) and its growing adoption by customers and farmers (Paull & Hennig, 2016) highlight the importance of considering how OF expansion may impact the dynamics of agricultural pests (e.g. Petit et al., 2020). Modeling approaches are useful tools to understand and forecast how pest densities and pest control may vary according to crop management and semi-natural habitats at the landscape scale (Begg et al. 2017). In this study, we modeled pest and predator abundances dynamics for contrasted scenarios of OF expansion in different landscapes. Our results indicate that, at the landscape scale, the IP (Isolated Plots converted first) scenario would provide the most benefits for conservation biological control (i.e. predator to pest ratio) in organic fields with little impact on pest densities in conventional fields.

### 1 Landscape context and OF expansion affect pest and predator densities

Populations responded to organic farming expansion in the landscape with up to a 437% increase and a 46% decrease in pest and predator densities, indicating that organic farming expansion could indeed lead to significant changes in biological control of pests in both organic and conventional fields depending on the landscape context. Predator densities generally increased or remained stable while pest densities either increased or decreased along OF expansion. When both pest and predator densities increased, predator densities increased more strongly than pest densities. The predator to pest ratio was about three to four times larger in organic than in conventional fields. Changes in pest and predator densities and their dynamics strongly depended on expansion scenarios in interaction with landscape contexts, i.e. the amount and fragmentation of SNH. Although most scenarios led to overall improvements in predator to pest ratios (seen here as a proxy of conservation biological control, CBC), some led to increases in pest densities, particularly in conventional fields which indicates that in some specific landscapes, carefully planning the spatial expansion of organic farming would be useful to avoid undesirable side effects.

From an ecological point of view, the predator to pest ratio dynamics observed in this study appeared driven by the dynamics of predators which was mostly dependent on the amount of semi-natural habitat (SNH). It was striking that only in landscapes with large proportion of SNH (SNH=50%), did predator densities increase very largely in organic fields and even increase slightly in conventional fields, leading to a decrease of pests in both types of fields. CBC also increased with landscape fragmentation in both OF and CF fields but mostly when the proportion of SNH was high. Since SNH fragmentation increased its edge length with cultivated habitats, this synergy between SNH amount and fragmentation on the level of CBC indicates the importance of predators’ spillover from semi-natural habitats on biological control. This interaction is also in line with the frequent observation that complex landscapes with more and more fragmented semi-natural habitats sustain more biodiversity within fields(Muneret, Auriol, Bonnard, et al.,2019; Smith et al., 2020; Tscharntke et al., 2021) and that reduced distance between SNH and crops favor spillover of predators (Holland et al., 2016; Jonsson et al., 2014; Lavandero et al., 2006; Tylianakis et al.,2006). Incidentally, it may indicate that landscape structures favoring the movement of predators into conventional fields may act as an ecological trap also luring non-target species to fields where mortality is high and long-term persistence impossible (Robertson & Hutto, 2006; Shelton & Badenes-Perez, 2006).Whatever the expansion scenario, we observed smooth changes in pest and predator densities. This was unexpected given the dramatic changes in the size of patches of OF and CF fields that underwent peculiar processes of progressive percolation/agglomeration (for OF) and its opposite disintegration (for CF) through the conversion of single central fields, leading to non-linear changes and even dramatic thresholds of mean OF and CF patch areas (especially in the IP scenario). This may indicate that the spatial smoothness of a conversion scenario over time is not necessary to maintain generally stable dynamics of biological control at landscape scale, neither in organic nor in conventional farms. This result is an indication that carefully curated temporal plans for OF expansion may not be not necessary, and that mean pest infestation risks may remain low for CF farms at the landscape scale. It contrasts with findings on the consequences of OF expansion in a pest - parasitoid model that indicated peaks of pest density for intermediate proportions of OF in the landscape (Bianchi et al. 2013). One main difference is that generalist predators, such as modeled here, may increase in density even in the absence of pests, thus limiting pest population peaks (Symondson et al., 2002).

### 2 One better scenario of organic farming expansion for CBC?

#### 2.1 A general pattern

The similar pest densities with all spatial expansion scenarios indicates that the choice of one scenario over another bears low risks, while potential benefits were more obvious with noticeable effects on predators. Both the level of conservation biological control (CBC) and pest densities have been used to evaluate the efficiency of pest control in spatial pest-predator models (Bianchi et al., 2013; Zamberletti et al., 2021). Yield or income have also been used (e.g. Le Gal et al., 2020; Milne et al., 2016). Here, because conventional fields relied on pesticides for pest control, and pest densities varied little in organic fields, CBC was a target mostly in organic fields while the main target for conventional fields was the density of pests.

Using these criteria, the IP scenario performed better, by improving CBC in organic fields and doing so at the expense of lower CBC, but not higher pest densities, in conventional fields. Regarding CBC, the IP scenario performed overall better for organic fields because of its clear positive effect on the predator to pest ratio. Patterns were more nuanced for conventional fields. While some scenario x landscape context combinations caused a small improvement in CBC, others caused strong decreases. Interestingly, the effect of expansion scenarios was weaker when they caused increases in CBC than when they caused decreases in CBC. Additionally, the ranking of scenarios was opposite in conventional vs organic fields (IP>RD>GP in organic fields vs GP>RD>IP in conventional fields). From the conventional farming point of view, the absence of planning (RD scenario) may thus constitute a reasonable scenario. However, pest densities in conventional fields were lower with IP than with the other scenarios. The best ranking of the IP scenario with higher CBC in organic fields and lower pest densities in conventional fields was observed in all landscape configurations, while some landscapes limited the decrease of CBC in conventional fields without canceling it.

The best performance of the IP scenario resulted from two distinct mechanisms: a predator spillover improving CBC in organic fields, and a combination of ‘chemical umbrella’ and lesser pest spillover in conventional fields. The IP scenario prioritized the conversion of fields neighboring organic fields or semi-natural areas. This meant that new organic fields benefitted from the spillover of predators from SNH, albeit with weak effects on pest density. Indeed, the predator to pest ratio improved mainly by an increase in predator numbers. Such trophic network top-heaviness can be caused by exogenous pathways that transfer energy into communities from across spatial and temporal boundaries: here, transfers from SNH (McCauley et al., 2018). With the IP scenario, CBC in organic fields may also have benefitted from a greater distance to conventional fields and from clustering of organic fields and SNH, as chemical pesticide use in conventional fields was modeled to kill most of the pests with strong side-effect on predators, drying both pest and predator populations in their surroundings through a sink effect. Clusters of conventional fields were more preserved from pests by the IP scenario, meaning that conventional fields benefitted from the protection of pesticides used in neighbor fields (the “chemical umbrella” effect) and that organic fields, a potential source of pests, were farther away from the conventional fields. A similar benefit of aggregating fields was found by Edwards et al. (2018) who simulated pest and predator dynamics in annual crops. Grouping annual crops could limit the abundance of dispersal limited pests because pests had to move over longer distances to reach new crop patches and reach central fields and could not build-up populations in the central fields. This best IP scenario is in accordance with the current trend of OF extension mostly happening in areas already rich in OF fields (Gabriel et al., 2009; Marton & Storm, 2021;Sánchez Herrera & Dimitri, 2019; Zollet & Maharjan, 2021).

#### 2.2 Effect of the landscape context on differences between scenarios

In our simulations, landscape configuration had a strong effect with differences in pest density up to two times for a given scenario. The proportion and fragmentation of SNH were generally of similar importance to the difference in the level of CBC between scenarios, although there was a clear decrease associated with the interaction between the two parameters, i.e. the difference between the IP and GP scenarios decreased with higher proportions and fragmentation of SNH. In conventional fields, this amounted mainly to the IP scenario that benefited slightly from SNH, while SNH did not affect pest density with the GP scenario. In organic fields, IP and GP converged at highest proportion and fragmentation levels, with pest densities of the GP scenario being favored while those of the IP scenario decreased. Interestingly, the IP scenario could bring higher benefits in organic fields in degraded landscapes, while both scenarios brought similar but lower benefits in preserved landscapes. This is consistent with the IP scenario breaking up large clusters of conventional fields, which were less present in landscapes with high proportions and fragmentation of SNH. Consequently, it may be less important to manage the OF expansion scenario in preserved landscapes, while the IP scenario should be favored in degraded landscapes.

#### 2.3 Robustness of the ranking of expansion scenarios

The ranking of OF expansion scenarios appeared robust to both the intensity and specificity of OF systems and the dispersal ability of pests and predators (Supplementary material S2). Varying these parameters did not affect the ranking of spatial expansion scenarios, only their relative differences. For example, intensive OF systems corresponding to intensive pest management (high pest mortality) were characterized by strong control of pest densities, therefore they showed little differences between scenarios. The only clear interaction between OF pest management and expansion scenario was in conventional fields: under extensive OF farming systems, the tendency of the IP scenario towards lower pest densities in conventional fields was reinforced (Supplementary material S2). This is because pest densities were overall higher for extensive OF systems but this did not strongly affect conventional fields, because, by limiting the decrease in CF patch size, the IP scenario resulted in less pest spill-over from organic to conventional fields. Further, dispersal ability had a marginal effect on pest densities (Supplementary material S3). Increasing dispersal tended to increase pest density’s response to landscape configuration (in particular to its fragmentation) in conventional fields, thus increasing differences in pest densities among expansion scenarios.

### 3 Limits and benefits of the modelling approach

Most mechanistic models of pest control by natural enemies are specific to a biological system and few address landscape scale crop management (reviewed in Alexandridis et al. (2021)). Such models generally comprise numerous parameters and allow deriving conclusions for specific landscape arrangements. Vinatier et al. (2012) for example showed that longer crop rotations reduced the parasitism of oilseed rape pollen beetle by decreasing the spatial and temporal connectivity of the resource for the parasitoid.

In the present study, we chose a mechanistic theoretical model that was based on few ecological processes and a very simplified representation of crop protection practices. Our focus was on comparing spatial scenarios and understanding how these interacted with the landscape patterns. It is recognized that the complexity of processes underlying conservation biological control in landscapes limits the ability of models to represent actual situations. For example, in reality many organisms show complex movement behavior (Gurarie et al., 2016) and interact within complex trophic networks, even in agricultural fields (Macfadyen et al., 2009). Further, while we assumed similar dispersal abilities for the pest and the predator, real species may have different dispersal abilities and thus perceive the landscape at a different grain (Jackson & Fahrig, 2012). Differences among scenarios would, for example, obviously be reduced for long-distance dispersers that would be less affected by landscape structure. We also made strong assumptions about the role of semi-natural habitats for pests and predators, assuming a generalist predator and a crop specialist pest that may survive in semi-natural habitats. Differences among spatial expansion scenarios would, for example, probably have been less if the pest had been able to reproduce in semi-natural habitats and would thus have been less sensitive to the spatial distribution of organic or conventional fields. Interestingly, despite these limitations, our conclusions about the best spatial scenario are consistent with those of the only pest-natural enemy spatially explicit model that, to our knowledge, addressed OF expansion (Bianchi, Ives and Schellhorn, 2013). Using a spatially explicit pest-parasitoid model these authors found that the spatial clustering of organic fields allowed a higher level of biocontrol in organic fields by protecting parasitoids from the detrimental effects of insecticides sprayed in conventional fields. In contrast to our results, however, they reported peaks of pests along OF expansion, possibly because, contrary to our assumptions, the parasitoid was specialized on the pest.

A last limitation of our approach is that results were averaged for organic and conventional fields at the landscape level. This simplification was driven by the large number of simulations to analyze. Aggregating outputs over space, however, may have masked local patterns and possible local peaks in pest densities. In a recent modelling study of a specialist pest and a generalist predator interacting in an heterogeneous agricultural landscape, Zamberletti et al. (2021, 2022) showed for example that semi-natural habitats increased the average landscape scale pest density (by reducing the number of necessary pesticide treatments) but locally reduced peaks of pest populations(Zamberletti et al., 2021, 2022). Further analyses of pest density dynamics at the field level would, thus, be necessary to confirm the better ranking of the IP scenario regarding local CBC and pest densities.

Despite these limitations, our approach set in light processes such as increased spill-over of predators in isolated fields, increased pest management efficiency in large patches of conventional fields and the importance of distance between organic and conventional fields, that help understand consequences of diverse organic farming expansion scenarios. They further highlight that landscape planning appeared most necessary when organic pest management had a low efficiency on pests and in landscapes with low quantities of semi-natural habitats.

## Conclusion

The scenario that consisted in setting the priority on isolated conventional fields for conversion to organic (IP) appeared as the most promising scenario to limit pest densities in conventional crops and improve CBC in organic crops, without increasing pest densities there. By examining a large number of landscape contexts and population parameters, we found that this result was robust but that landscape planning appeared most necessary when organic pesticides had a low efficiency on pests. Furthermore, landscape contexts with large proportions and fragmentation of semi-natural habitats supported the highest level of CBC. The modeling of agricultural landscapes is still a research objective (Poggi et al., 2018) and improving both the consideration of agricultural practices and the calibration of models using observed data regarding the life history traits of pests and predators will hopefully help design agroecological landscapes.

## Acknowledgements

The work was partially supported by the PEERLESS project (ANR-12-AGRO-0006). The authors thank Lionel Roques and Mamadou Ciss for valuable discussions at the genesis of the work.

## Data and code availability

The model used in this study is based on a model developed by Martinet and Roques (2022) that is available on the following public repository: https://doi.org/10.17605/OSF.IO/Z2QCX

The model outputs of the present study, as well as the R scripts used to build graphs and analyse data are available on the Zenodo public repository: https://doi.org/10.5281/zenodo.6597282

## Supplementary materials

### SM1 Effects of semi-natural habitat fragmentation and OF expansion scenarios on landscape structure

#### Landscape structure

Increasing semi-natural habitat (SNH) fragmentation (parameter *fr*) resulted in an increase in the number of patches of each habitat type (SNH but also organic farming (OF) and conventional farming (CF)) (Figure S1.1) as well as an increase in edge length among habitat type (Figure S1.2). Patch area and edge length were calculated with R package landscapemetrics (Hesselbarth et al. 2019, Ecography 42:1648-1657)

**Figure S1.1.**
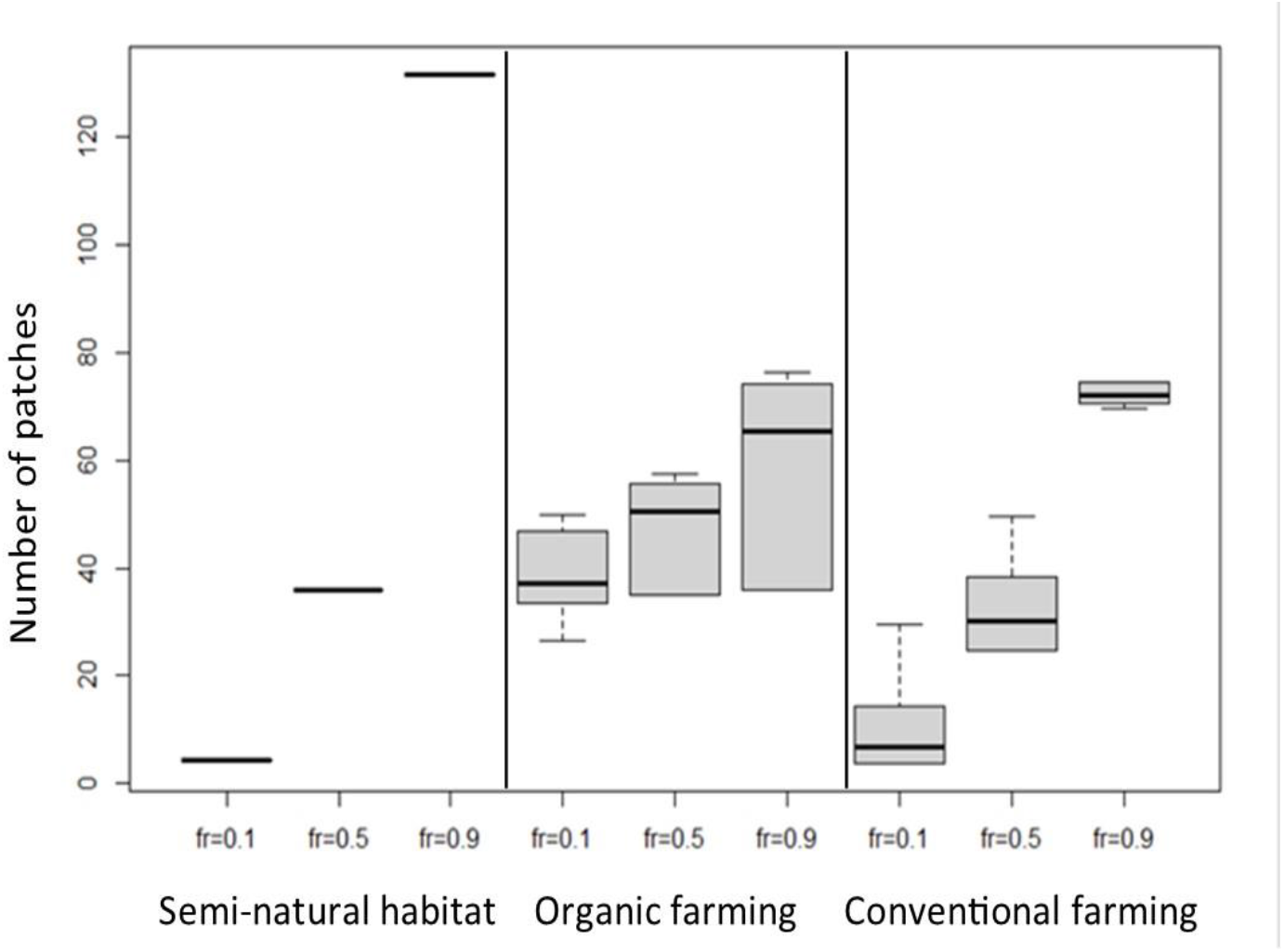
Number of patches per landscape and habitat type for the three levels of semi-natural habitat fragmentation fr. The graph presents box plots of all values pooled over the 11664 landscapes at the end of simulations of OF expansion (t=50) The dark line is the median, whiskers represent the first and third quartiles.

**Figure S1.2.**
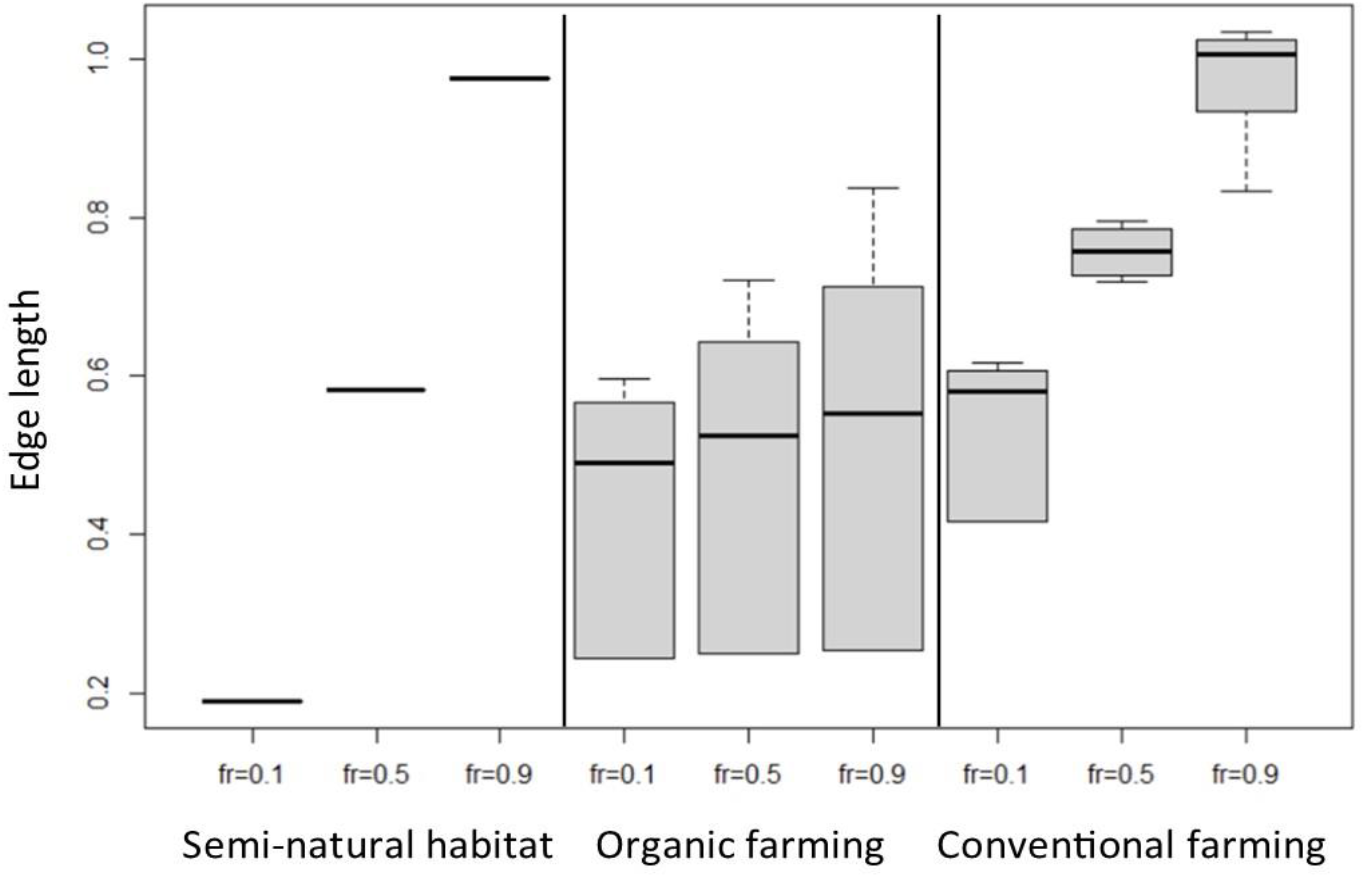
Edge length (in pixel side length per pixel) per landscape by habitat type for the three levels of semi-natural habitat fragmentation *fr*. The graph presents box plots of all values pooled over the 11664 landscapes at the end of simulations of OF expansion (t=50). The dark line is the median, whiskers represent the first and third quartiles.

#### Dynamics of the number and area of organic and conventional patches

As expected, changes in the areas and numbers of organic and conventional patches along organic farming expansion depended on the landscape characteristics (amount of semi-natural habitat and its fragmentation) and on the organic farming expansion scenario. Organic and conventional patches were overall larger and less numerous in landscapes where the amount of semi-natural habitat was small and little fragmented (upper left Fig. S1.3) indicating in particular that the level of semi-natural habitat fragmentation translated to overall landscape fragmentation.

Overall the dynamics of the patch area were driven by two processes. Indeed, the conversion of individual fields from conventional to organic may lead to progressive changes in conventional patch area, either increasing it when converted fields were isolated and/or decreasing it when converted fields were part of a larger patch. In this second situation, the conversion of a single conventional field may occasionally lead to the splitting of a large conventional patch. Such splitting led to large drops in the mean conventional patch area (eg. Fig.S1.3, at 35 years for the GP scenario with fr=0.1 and SNH=10%). The symmetrical process of merging organic patches following the conversion of individual fields may create a sudden large increase in mean organic patch area. This last process occurred when the organic share was high enough over the landscape.

**Figure S1.3.**
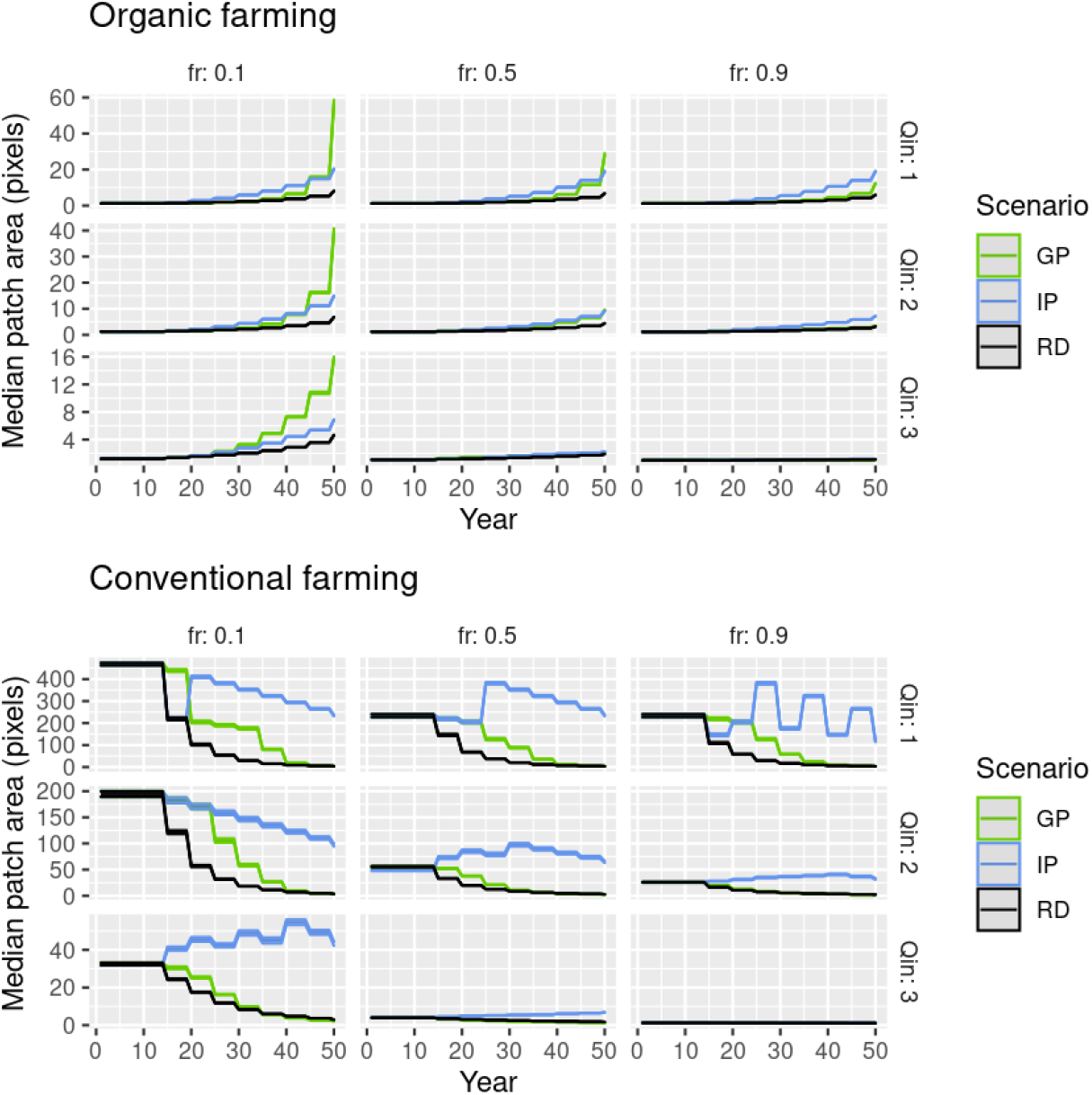
Effects of OF expansion scenarios on the area of organic and conventional patches. Landscape change (median patch area in pixels) during the simulations. The envelope around each curve represents the standard error.

Because they set different priorities regarding field conversion, the different scenarios led to different mean patch area dynamics. Constraints were furthermore imposed by the spatial distribution of semi-natural areas. The IP scenario always resulted in conventional patches that were larger than the other scenarios. This is because, when available, conventional fields in the smallest conventional patches were converted to organic which resulted in an initial disappearance of small conventional patches and thus an increase in average conventional patch area. When these small patches were all converted, larger ones started being partially converted to organic, leading to a secondary decrease in conventional patch area (Fig S1.3, after ca. 25 years). These two trends (increase then decrease) were observed in landscapes with both small and large conventional patches initially, i.e. moderately fragmented landscapes with a small to moderate proportion of semi-natural habitat. In little fragmented landscapes with few semi-natural habitats (upper left panel, Fig. S1.3) all conventional patches were large initially so that patch size decreased slowly from the beginning of organic farming expansion. In contrast, in highly fragmented landscapes with a high proportion of semi-natural habitat (lower right panel, Fig. S1.3), there were mostly isolated conventional fields initially so that patch size remained almost constant. The GP scenario, by eroding small parts of large conventional patches at first, slowly and moderately reduced the average conventional patch area. This decrease accelerated in a second step when the erosion incidentally led to the splitting of the still rather large conventional patches into smaller ones. This process was strongest in landscapes with large conventional patches initially, i.e. little fragmented or with a small proportion of semi-natural habitats (left column and upper row panels, Fig. S1.3). Lastly, the RD scenario led to a progressive reduction of conventional patch area by both converting fields located in small patches and reducing the area of large conventional patches.

The effect of organic expansion on the area and number of organic patches was consistent with the above changes to conventional patches. Whatever the expansion scenario, when the landscape was very fragmented and with a large proportion of semi-natural habitat, conversion of conventional fields increased the number of organic fields but not their average area, conventional patches being mostly composed of single fields (lower right panel, Fig.S1.3). Mean organic patch area increased in all other situations.

Mean organic patch area increased most at first with the IP scenario, particularly when the landscape was a little fragmented (left column, Fig. S1.3) because conventional fields that were converted tended to be neighboring already organic fields. In contrast, with the GP scenario, organic fields first tended to be isolated from other organic fields so that the average patch area increased slowly. However, when the landscape was little fragmented (left column, Fig. S1.3), these small organic patches merged when the proportion of organic farming increased and the average organic patch area increased sharply while the number of patches decreased.

### SM2 Effect of the type of organic farming on pest densities and interaction with expansion scenario

Pests were on average more abundant in both types of fields when organic farming was less intensive, i.e. pest management affected pest population growth less (Table 3) and, to a lesser extent, when it was less specific, i.e. there was a small differential in pest management-induced mortality between predators and pests (Table 3). In organic fields, the intensity of organic farming affected pest abundance far more than specificity, regardless of the amount of semi-natural habitat and its level of fragmentation.

As expected, the effect of organic farming intensity and specificity was much less pronounced in conventional fields. The effect of specificity was very weak. The effect of OF intensity was observable mainly in landscapes that were characterized by a low fragmentation (figure S2) Interestingly, the response of pest density to expansion scenario showed the same pattern whatever the OF type. It was very similar whatever the expansion scenario in organic fields and pest densities were generally lower for the IP scenario in conventional fields.

The fact that pest management specificity generally had little effect except for the extensive OF systems, confirmed the low effect of predators on pest densities in conventional fields, and the high impact of pest management compared to CBC in our simulations. For extensive organic systems, organic fields were possibly a source of pests for surrounding fields. Indeed, we observed more pests in conventional fields when organic farming systems were extensive, possibly indicating pest spillover from OF fields with higher pest populations. The latter is supported by the fact that the effect of OF farming system on pest density in CF was reduced in some landscape configurations. Specifically, conventional fields in landscapes with high proportion of SNH were less sensitive to OF farming system intensity, possibly because of lesser proximity to OF sources, and because of higher predator’s spillover from SNH.

**Figure S2.**
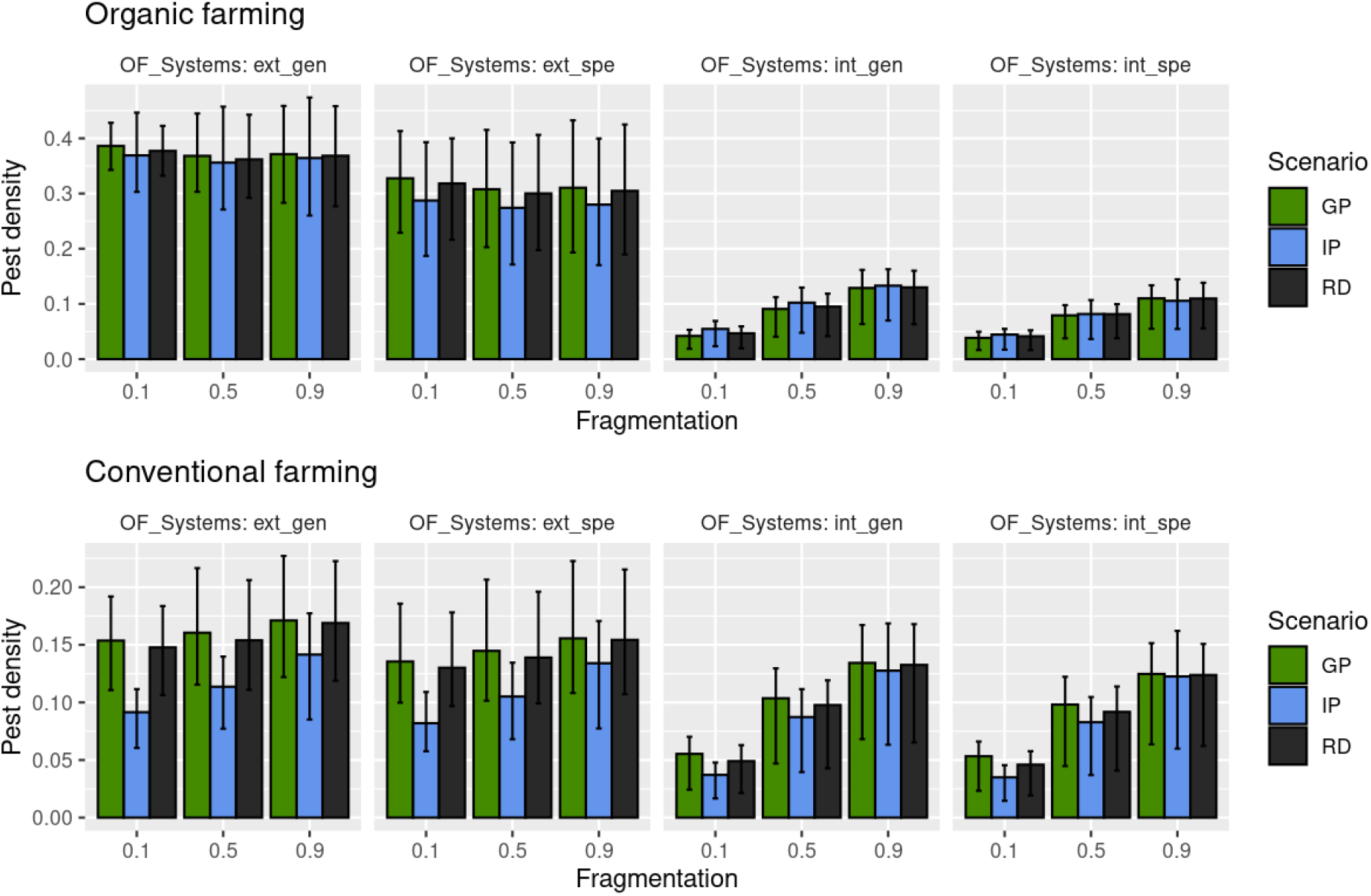
Effects of organic farming expansion scenario, organic farming type and landscape structure on the density of pests in organic and conventional fields. “ext” and “int”: low vs high pest management intensity, respectively. “spe” vs “gen”: specific vs generalist pest management practices, respectively (see Table. 3). Error bars represent standard deviations over landscapes.

### SM3 Effect of pests dispersal and SNH fragmentation on pest densities and interaction with OF expansion scenario

Pest dispersal had a lower effect than the other parameters with a maximum delta of ±0.05 in pest densities (Fig.S3). Pest densities in organic and conventional fields were overall higher when pest dispersal was high but this effect was weak, and mainly observable in conventional fields. In both types of fields, the positive effect of dispersal increased with the level of fragmentation of semi-natural habitats (for example, in conventional fields, for the GP scenario, pest density increased by 0.01 when *fr*=0.1, and by 0.04 when *fr*=0.9 – Fig. S3). There was one exception to this trend with a small decrease in pest density with dispersal. It was observed with the IP scenario in organic fields (from 0.19 to 0.18 for *fr*=0.1, Fig. S3).

The increase in densities with dispersal, fragmentation and their interaction was probably due to a higher ability of pests to avoid CBC-heavy areas (near SNH, which are sources of predators) and to reach resource-rich areas. Globally speaking, dispersal ability amplified the effect of every landscape parameter (fragmentation, expansion scenario).

**Figure S3.**
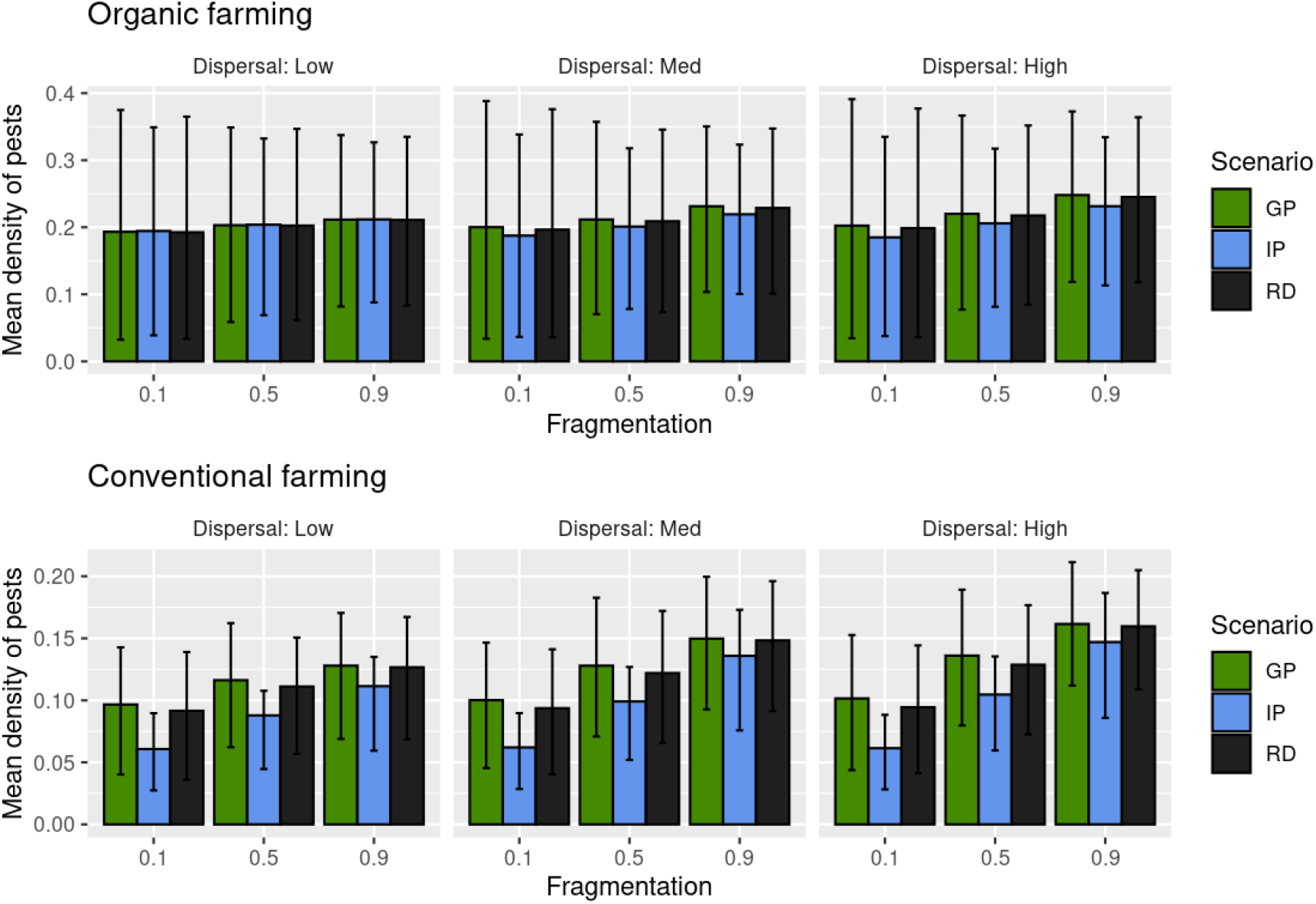
Mean density of pests as a function of pest dispersal, semi-natural habitat fragmentation and organic farming expansion scenario

## Notes

https://doi.org/10.17605/OSF.IO/Z2QCX

https://doi.org/10.5281/zenodo.7576542

### Competing Interest Statement

The authors have declared no competing interest.

### Summary of Updates

This manuscript is the result of a first round of reviews PCI Ecology's editorial board and reviewers. We took into account all their comments, which can be found with our answers on PCI Ecology's website. Changes in the manuscript ar highlighted in blue. We considerably refocused the manuscript by moving to supplementary material every result analysis that did not specifically address the comparison of expansion scenarios. We also added a general figure explaining the simulation study and numerous details about the population dynamics modelling and its relation to the real-world ecology of pests and predators. Please also note that the data repository has been updated to take into account some of the comments, and can now be found here: https://doi.org/10.5281/zenodo.7576542

## References

Adl, S., Iron, D., & Kolokolnikov, T. (2011). A threshold area ratio of organic to conventional agriculture causes recurrent pathogen outbreaks in organic agriculture. Science of The Total Environment, 409(11), 2192–2197. https://doi.org/10.1016/j.scitotenv.2011.02.026

Alexandridis, N., Marion, G., Chaplin-Kramer, R., Dainese, M., Ekroos, J., Grab, H., Jonsson, M., Karp, D. S., Meyer, C., O’Rourke, M. E., Pontarp, M., Poveda, K., Seppelt, R., Smith, H. G., Martin, E. A., & Clough, Y. (2021). Models of natural pest control: Towards predictions across agricultural landscapes. Biological Control, 163, 104761. https://doi.org/10.1016/j.biocontrol.2021.104761

Batáry, P., Báldi, A., Kleijn, D., & Tscharntke, T. (2011). Landscape-moderated biodiversity effects of agri-environmental management: A meta-analysis. Proc. R. Soc. B., 1894–1902.

Begg, G. S., Cook, S. M., Dye, R., Ferrante, M., Franck, P., Lavigne, C., Lövei, G. L., Mansion-Vaquie, A., Pell, J. K., Petit, S., Quesada, N., Ricci, B., Wratten, S. D., & Birch, A. N. E. (2017). A functional overview of conservation biological control. Crop Protection, 97, 145–158. https://doi.org/10.1016/j.cropro.2016.11.008

Bianchi, F. J. J. A., Ives, A. R., & Schellhorn, N. A. (2013). Interactions between conventional and organic farming for biocontrol services across the landscape. Ecological Applications, 23(7), 1531–1543. https://doi.org/10.1890/12-1819.1

Caprio, E., Nervo, B., Isaia, M., Allegro, G., & Rolando, A. (2015). Organic versus conventional systems in viticulture: Comparative effects on spiders and carabids in vineyards and adjacent forests. Agricultural Systems, 136, 61–69. https://doi.org/10.1016/j.agsy.2015.02.009

Chaplin-Kramer, R., O’Rourke, M. E., Blitzer, E. J., & Kremen, C. (2011). A meta-analysis of crop pest and natural enemy response to landscape complexity. Ecology Letters, 14(9), 922–932. https://doi.org/10.1111/j.1461-0248.2011.01642.x

Ciss, M., Poggi, S., Memmah, M., Franck, P., Gosme, M., Parisey, N., & Roques, L. (2016). A model-based approach to assess the effectiveness of pest biocontrol by natural enemies (p. np). auto-saisine. https://hal.archives-ouvertes.fr/hal-01609595

Colbach, N., Petit, S., Chauvel, B., Deytieux, V., Lechenet, M., Munier-Jolain, N., & Cordeau, S. (2020). The Pitfalls of Relating Weeds, Herbicide Use, and Crop Yield: Don’t Fall Into the Trap ! A Critical Review. Frontiers in Agronomy, 2, 33. https://doi.org/10.3389/fagro.2020.615470

Concepción, E. D., Díaz, M., & Baquero, R. A. (2008). Effects of landscape complexity on the ecological effectiveness of agri-environment schemes. Landscape Ecology, 23(2), 135–148. https://doi.org/10.1007/s10980-007-9150-2

Deshayes, C., Siegwart, M., Pauron, D., Froger, J.-A., Lapied, B., & Apaire-Marchais, V. (2017). Microbial Pest Control Agents: Are they a Specific And Safe Tool for Insect Pest Management? Current Medicinal Chemistry, 24(27), 2959–2973. https://doi.org/10.2174/0929867324666170314144311

Dib, H., Sauphanor, B., & Capowiez, Y. (2010). Effect of codling moth exclusion nets on the rosy apple aphid, Dysaphis plantaginea, and its control by natural enemies. Crop Protection, 29(12), 1502–1513. https://doi.org/10.1016/j.cropro.2010.08.012

Dib, H., Sauphanor, B., & Capowiez, Y. (2016). Effect of management strategies on arthropod communities in the colonies of rosy apple aphid, Dysaphis plantaginea Passerini (Hemiptera: Aphididae) in south-eastern France. Agriculture, Ecosystems & Environment, 216, 203–206. https://doi.org/10.1016/j.agee.2015.10.003

Diekötter, T., Wamser, S., Dörner, T., Wolters, V., & Birkhofer, K. (2016). Organic farming affects the potential of a granivorous carabid beetle to control arable weeds at local and landscape scales. Agricultural and Forest Entomology, 18(2), 167–173. Scopus. https://doi.org/10.1111/afe.12150

Diekötter, T., Wamser, S., Wolters, V., & Birkhofer, K. (2010). Landscape and management effects on structure and function of soil arthropod communities in winter wheat. Special section Harvested perennial grasslands: Ecological models for farming’s perennial future, 137(1), 108–112. https://doi.org/10.1016/j.agee.2010.01.008

Djoudi, E. A., Marie, A., Mangenot, A., Puech, C., Aviron, S., Plantegenest, M., & Pétillon, J. (2018). Farming system and landscape characteristics differentially affect two dominant taxa of predatory arthropods. Agriculture, Ecosystems & Environment, 259, 98–110. https://doi.org/10.1016/j.agee.2018.02.031

Djoudi, E. A., Plantegenest, M., Aviron, S., & Pétillon, J. (2019). Local vs. Landscape characteristics differentially shape emerging and circulating assemblages of carabid beetles in agroecosystems. Agriculture, Ecosystems & Environment, 270-271, 149–158. https://doi.org/10.1016/j.agee.2018.10.022

Duru, M., Therond, O., Martin, G., Martin-Clouaire, R., Magne, M.-A., Justes, E., Journet, E.-P., Aubertot, J.-N., Savary, S., Bergez, J.-E., & Sarthou, J. P. (2015). How to implement biodiversity-based agriculture to enhance ecosystem services: A review. Agronomy for Sustainable Development, 35(4), 1259–1281. https://doi.org/10.1007/s13593-015-0306-1

Edwards C.B., Rosenheim J.A., & Segoli M. (2018) Aggregating fields of annual crops to form larger-scale monocultures can suppress dispersal-limited herbivores. Theoretical Ecology, 11 (3), 321–331 https://doi.org/10.1007/s12080-018-0369-0

Fahrig, L. (2003). Effects of Habitat Fragmentation on Biodiversity. In Annual Review of Ecology, Evolution, and Systematics (Vol. 34, Numéro 1, p. 487–515).

Fahrig, L., & Paloheimo, J. (1988). Effect of Spatial Arrangement of Habitat Patches on Local Population Size. Ecology, 69(2), 468–475. https://doi.org/10.2307/1940445

Gabriel, D., Carver, S. J., Durham, H., Kunin, W. E., Palmer, R. C., Sait, S. M., Stagl, S., & Benton, T. G. (2009). The spatial aggregation of organic farming in England and its underlying environmental correlates. Journal of Applied Ecology, 46(2), 323–333. https://doi.org/10.1111/j.1365-2664.2009.01624.x

Gomiero, T. (2018). Food quality assessment in organic vs. Conventional agricultural produce: Findings and issues. HUMUSICA 3-Reviews, Applications, Tools, 123, 714–728. https://doi.org/10.1016/j.apsoil.2017.10.014

Gosme, M., de Villemandy, M., Bazot, M., & Jeuffroy, M.-H. (2012). Local and neighbourhood effects of organic and conventional wheat management on aphids, weeds, and foliar diseases. Agriculture, Ecosystems & Environment, 161, 121–129. https://doi.org/10.1016/j.agee.2012.07.009

Goulson Dave, Nicholls Elizabeth, Botías Cristina, & Rotheray Ellen L. (2015). Bee declines driven by combined stress from parasites, pesticides, and lack of flowers. Science, 347(6229), 1255957. https://doi.org/10.1126/science.1255957

Graillot, B., Bayle, S., Blachere-Lopez, C., Besse, S., Siegwart, M., & Lopez-Ferber, M. (2016). Biological Characteristics of Experimental Genotype Mixtures of Cydia Pomonella Granulovirus (CpGV): Ability to Control Susceptible and Resistant Pest Populations. Viruses, 8(5). https://doi.org/10.3390/v8050147

Gurarie, E., Bracis, C., Delgado, M., Meckley, T. D., Kojola, I., & Wagner, C. M. (2016). What is the animal doing ? Tools for exploring behavioural structure in animal movements. Journal of Animal Ecology, 85(1), 69–84. https://doi.org/10.1111/1365-2656.12379

Heimpel, G. E., & Mills, N. J. (2017). Biological Control: Ecology and Applications. Cambridge University Press; Cambridge Core. https://doi.org/10.1017/9781139029117

Hillaert, J., Vandegehuchte, M., Hovestadt, T., & Bonte, D. (2018). Information use during movement regulates how fragmentation and loss of habitat affect body size. https://doi.org/10.1101/265025

Hillaert, J., Vandegehuchte, M. L., Hovestadt, T., & Bonte, D. (2020). Habitat loss and fragmentation increase realized predator–prey body size ratios. Functional Ecology, 34(2), 534–544. https://doi.org/10.1111/1365-2435.13472

Holland, J. M., Bianchi, F. J., Entling, M. H., Moonen, A.-C., Smith, B. M., & Jeanneret, P. (2016). Structure, function and management of semi-natural habitats for conservation biological control: A review of European studies. Pest Management Science, 72(9), 1638–1651. https://doi.org/10.1002/ps.4318

Ilbery, B., Holloway, L., & Arber, R. (1999). The Geography of Organic Farming in England and Wales in the 1990s. Tijdschrift voor Economische en Sociale Geografie, 90(3), 285–295. https://doi.org/10.1111/1467-9663.00070

Inclán, D. J., Cerretti, P., Gabriel, D., Benton, T. G., Sait, S. M., Kunin, W. E., Gillespie, M. A. K., & Marini, L. (2015). Organic farming enhances parasitoid diversity at the local and landscape scales. Journal of Applied Ecology, 52(4), 1102–1109. https://doi.org/10.1111/1365-2664.12457

Jackson, H. B., & Fahrig, L. (2012). What size is a biologically relevant landscape? Landscape Ecology, 27(7), 929–941. https://doi.org/10.1007/s10980-012-9757-9

Jonsson, M., Bommarco, R., Ekbom, B., Smith, H. G., Bengtsson, J., Caballero-Lopez, B., Winqvist, C., & Olsson, O. (2014). Ecological production functions for biological control services in agricultural landscapes. Methods in Ecology and Evolution, 5(3), 243–252. https://doi.org/10.1111/2041-210X.12149

Juhel, A. S., Barbu, C. M., Franck, P., Roger-Estrade, J., Butier, A., Bazot, M., & Valantin-Morison, M. (2017). Characterization of the pollen beetle, Brassicogethes aeneus, dispersal from woodlands to winter oilseed rape fields. PLOS ONE, 12(8), e0183878. https://doi.org/10.1371/journal.pone.0183878

Karp, D. S., Chaplin-Kramer, R., Meehan, T. D., Martin, E. A., DeClerck, F., Grab, H., Gratton, C., Hunt, L., Larsen, A. E., Martínez-Salinas, A., O’Rourke, M. E., Rusch, A., Poveda, K., Jonsson, M., Rosenheim, J. A., Schellhorn, N. A., Tscharntke, T., Wratten, S. D., Zhang, W.,… Zou, Y. (2018). Crop pests and predators exhibit inconsistent responses to surrounding landscape composition. Proceedings of the National Academy of Sciences, 115(33), E7863. https://doi.org/10.1073/pnas.1800042115

Knapp, S., & van der Heijden, M. G. A. (2018). A global meta-analysis of yield stability in organic and conservation agriculture. Nature Communications, 9(1), 3632. https://doi.org/10.1038/s41467-018-05956-1

Kremen, C., Williams, N. M., Aizen, M. A., Gemmill-Herren, B., LeBuhn, G., Minckley, R., Packer, L., Potts, S. G., Roulston, T., Steffan-Dewenter, I., Vázquez, D. P., Winfree, R., Adams, L., Crone, E. E., Greenleaf, S. S., Keitt, T. H., Klein, A.-M., Regetz, J., & Ricketts, T. H. (2007). Pollination and other ecosystem services produced by mobile organisms: A conceptual framework for the effects of land-use change. Ecology Letters, 10(4), 299–314. https://doi.org/10.1111/j.1461-0248.2007.01018.x

Lavandero, B., Wratten, S. D., Didham, R. K., & Gurr, G. (2006). Increasing floral diversity for selective enhancement of biological control agents: A double-edged sward? Basic and Applied Ecology, 7(3), 236–243. https://doi.org/10.1016/j.baae.2005.09.004

Le Gal, A., Robert, C., Accatino, F., Claessen, D., & Lecomte, J. (2020). Modelling the interactions between landscape structure and spatio-temporal dynamics of pest natural enemies: Implications for conservation biological control. Ecological Modelling, 420, 108912. https://doi.org/10.1016/j.ecolmodel.2019.108912

Lefebvre, M., Franck, P., Toubon, J.-F., Bouvier, J.-C., & Lavigne, C. (2016). The impact of landscape composition on the occurrence of a canopy dwelling spider depends on orchard management. Agriculture, Ecosystems & Environment, 215, 20–29. https://doi.org/10.1016/j.agee.2015.09.003

Lourenço, R., Pereira, P. F., Oliveira, A., Ribeiro-Silva, J., Figueiredo, D., Rabaça, J. E., Mira, A., & Marques, J. T. (2021). Effect of vineyard characteristics on the functional diversity of insectivorous birds as indicator of potential biocontrol services. Ecological Indicators, 122, 107251. https://doi.org/10.1016/j.ecolind.2020.107251

Macfadyen, S., Gibson, R., Polaszek, A., Morris, R. J., Craze, P. G., Planqué, R., Symondson, W. O. C., & Memmott, J. (2009). Do differences in food web structure between organic and conventional farms affect the ecosystem service of pest control? Ecology Letters, 12(3), 229–238. https://doi.org/10.1111/j.1461-0248.2008.01279.x\uc0\u160{}

Marliac, G., Penvern, S., Barbier, J.-M., Lescourret, F., & Capowiez, Y. (2015). Impact of crop protection strategies on natural enemies in organic apple production. Agronomy for Sustainable Development, 35(2), 803–813. https://doi.org/10.1007/s13593-015-0282-5

Martinet, V., & Roques, L. (2022). An ecological-economic model of land-use decisions, agricultural production and biocontrol. R. Soc. open sci. 9:220169. https://doi.org/10.1098/rsos.220169

Marton, T. A., & Storm, H. (2021). The case of organic dairy conversion in Norway: Assessment of multivariate neighbourhood effects. Q Open, 1(1), qoab009. https://doi.org/10.1093/qopen/qoab009

MATLAB. (2018a). The MathWorks, Inc.

McCauley, D. J., Gellner, G., Martinez, N. D., Williams, R. J., Sandin, S. A., Micheli, F., Mumby, P. J., & McCann, K. S. (2018). On the prevalence and dynamics of inverted trophic pyramids and otherwise top-heavy communities. Ecology Letters, 21(3), 439–454. https://doi.org/10.1111/ele.12900

Milne, A. E., Bell, J. R., Hutchison, W. D., van den Bosch, F., Mitchell, P. D., Crowder, D., Parnell, S., & Whitmore, A. P. (2016). The Effect of Farmers’ Decisions on Pest Control with Bt Crops: A Billion Dollar Game of Strategy. PLOS Computational Biology, 11(12), e1004483. https://doi.org/10.1371/journal.pcbi.1004483

Mózner, Z., Tabi, A., & Csutora, M. (2012). Modifying the yield factor based on more efficient use of fertilizer—The environmental impacts of intensive and extensive agricultural practices. Ecological Indicators, 16, 58–66. https://doi.org/10.1016/j.ecolind.2011.06.034

Muneret, L., Auriol, A., Bonnard, O., Richart-Cervera, S., Thiéry, D., & Rusch, A. (2019). Organic farming expansion drives natural enemy abundance but not diversity in vineyard-dominated landscapes. Ecology and Evolution, 9(23), 13532–13542. https://doi.org/10.1002/ece3.5810

Muneret, L., Auriol, A., Thiéry, D., & Rusch, A. (2019). Organic farming at local and landscape scales fosters biological pest control in vineyards. Ecological Applications, 29(1), e01818. https://doi.org/10.1002/eap.1818

Muneret, L., Thiéry, D., Joubard, B., & Rusch, A. (2018). Deployment of organic farming at a landscape scale maintains low pest infestation and high crop productivity levels in vineyards. Journal of Applied Ecology, 55(3), 1516–1525. https://doi.org/10.1111/1365-2664.13034

Muth, F., & Leonard, A. S. (2019). A neonicotinoid pesticide impairs foraging, but not learning, in free-flying bumblebees. Scientific Reports, 9(1), 4764. https://doi.org/10.1038/s41598-019-39701-5

ORAB PACA. (2020). Les chiffres clés de ?agriculture biologique en PACA. https://www.bio-provence.org/Chiffres-cles-de-la-bio-en-PACA-120

Pärn, J., Pinay, G., & Mander, Ü. (2012). Indicators of nutrients transport from agricultural catchments under temperate climate: A review. Adaptation and functional water management by land use change, 22, 4–15. https://doi.org/10.1016/j.ecolind.2011.10.002

Paull, J., & Hennig, B. D. (2016). Atlas of Organics: Four maps of the world of organic agriculture.

Perez-Alvarez, R., Nault, B. A., & Poveda, K. (2019). Effectiveness of augmentative biological control depends on landscape context. Scientific Reports, 9(1), 8664. https://doi.org/10.1038/s41598-019-45041-1

Petit, S., Muneret, L., Carbonne, B., Hannachi, M., Ricci, B., Rusch, A., & Lavigne, C. (2020). Chapter One—Landscape-scale expansion of agroecology to enhance natural pest control: A systematic review. In D. A. Bohan & A. J. Vanbergen (Éds.), Advances in Ecological Research (Vol. 63, p. 1–48). Academic Press. https://doi.org/10.1016/bs.aecr.2020.09.001

Poggi, S., Papaïx, J., Lavigne, C., Angevin, F., Le Ber, F., Parisey, N., Ricci, B., Vinatier, F., & Wohlfahrt, J. (2018). Issues and challenges in landscape models for agriculture: From the representation of agroecosystems to the design of management strategies. Landscape Ecology, 33(10), 1679–1690. https://doi.org/10.1007/s10980-018-0699-8

Puech, C., Poggi, S., Baudry, J., & Aviron, S. (2015). Do farming practices affect natural enemies at the landscape scale? Landscape Ecology, 30(1), 125–140. https://doi.org/10.1007/s10980-014-0103-2

R Software (3.5.2). (2017). [Computer software]. Development Core Team.

Ricci, B., Franck, P., Toubon, J. F., Bouvier, J. C., Sauphanor, B., & Lavigne, C. (2009). The influence of landscape on insect pest dynamics: A case study in southeastern France. Landscape Ecology, 24(3), 337–349. https://doi.org/10.1007/s10980-008-9308-6

Ricci, B., Lavigne, C., Alignier, A., Biju-Duval, L., Bouvier, J., Choisis, J.-P., Franck, P., Joannon, A., Ladet, S., Mezerette, F., Plantegenest, M., Savary, G., Thomas, C., Vialatte, A., & Petit, S. (2019). Local pesticide use intensity conditions landscape effects on biological pest control. Proceedings. Biological sciences, 286, 20182898. https://doi.org/10.1098/rspb.2018.2898

Robertson, B. A., & Hutto, R. L. (2006). A FRAMEWORK FOR UNDERSTANDING ECOLOGICAL TRAPS AND AN EVALUATION OF EXISTING EVIDENCE. Ecology, 87(5), 1075–1085. https://doi.org/10.1890/0012-9658(2006)87[1075:AFFUET]2.0.CO;2

Roques, L. (2015). MULTILAND: a neutral landscape generator designed for theoretical studies.

Salomé, M., Kesse-Guyot, E., Fouillet, H., Touvier, M., Hercberg, S., Huneau, J.-F., & Mariotti, F. (2021). Development and evaluation of a new dietary index assessing nutrient security by aggregating probabilistic estimates of the risk of nutrient deficiency in two French adult populations. British Journal of Nutrition, 126(8), 1225–1236. Cambridge Core. https://doi.org/10.1017/S0007114520005115

Sánchez Herrera, J. C., & Dimitri, C. (2019). The Role of Clustering in the Adoption of Organic Dairy: A Longitudinal Networks Analysis between 2002 and 2015. Sustainability, 11(6). https://doi.org/10.3390/su11061514

Sánchez-Bayo, F., & Wyckhuys, K. A. G. (2019). Worldwide decline of the entomofauna: A review of its drivers. Biological Conservation, 232, 8–27. https://doi.org/10.1016/j.biocon.2019.01.020

Shelton, A. M., & Badenes-Perez, F. R. (2006). CONCEPTS AND APPLICATIONS OF TRAP CROPPING IN PEST MANAGEMENT. Annual Review of Entomology, 51(1), 285–308. https://doi.org/10.1146/annurev.ento.51.110104.150959

Sirami, C., Gross, N., Baillod, A. B., Bertrand, C., Carrié, R., Hass, A., Henckel, L., Miguet, P., Vuillot, C., Alignier, A., Girard, J., Batáry, P., Clough, Y., Violle, C., Giralt, D., Bota, G., Badenhausser, I., Lefebvre, G., Gauffre, B.,… Fahrig, L. (2019). Increasing crop heterogeneity enhances multitrophic diversity across agricultural regions. Proceedings of the National Academy of Sciences, 116(33), 16442. https://doi.org/10.1073/pnas.1906419116

Smith, O. M., Cohen, A. L., Reganold, J. P., Jones, M. S., Orpet, R. J., Taylor, J. M., Thurman, J. H., Cornell, K. A., Olsson, R. L., Ge, Y., Kennedy, C. M., & Crowder, D. W. (2020). Landscape context affects the sustainability of organic farming systems. Proceedings of the National Academy of Sciences, 201906909. https://doi.org/10.1073/pnas.1906909117

Symondson, W. O. C., Sunderland, K. D., & Greenstone, M. H. (2002). Can Generalist Predators be Effective Biocontrol Agents? Annual Review of Entomology, 47(1), 561–594. https://doi.org/10.1146/annurev.ento.47.091201.145240

Tscharntke, T., Grass, I., Wanger, T. C., Westphal, C., & Batáry, P. (2021). Beyond organic farming – harnessing biodiversity-friendly landscapes. Trends in Ecology & Evolution, 36(10), 919–930. https://doi.org/10.1016/j.tree.2021.06.010

Tscharntke, T., Karp, D. S., Chaplin-Kramer, R., Batáry, P., DeClerck, F., Gratton, C., Hunt, L., Ives, A., Jonsson, M., Larsen, A., Martin, E. A., Martínez-Salinas, A., Meehan, T. D., O’Rourke, M., Poveda, K., Rosenheim, J. A., Rusch, A., Schellhorn, N., Wanger, T. C.,… Zhang, W. (2016). When natural habitat fails to enhance biological pest control – Five hypotheses. Biological Conservation, 204, 449–458. https://doi.org/10.1016/j.biocon.2016.10.001

Tscharntke, T., Klein, A. M., Kruess, A., Steffan-Dewenter, I., & Thies, C. (2005). Landscape perspectives on agricultural intensification and biodiversity – ecosystem service management. Ecology Letters, 8(8), 857–874. https://doi.org/10.1111/j.1461-0248.2005.00782.x

Tscharntke, T., Tylianakis, J. M., Rand, T. A., Didham, R. K., Fahrig, L., Batáry, P., Bengtsson, J., Clough, Y., Crist, T. O., Dormann, C. F., Ewers, R. M., Fründ, J., Holt, R. D., Holzschuh, A., Klein, A. M., Kleijn, D., Kremen, C., Landis, D. A., Laurance, W.,… Westphal, C. (2012). Landscape moderation of biodiversity patterns and processes—Eight hypotheses. Biological Reviews, 87(3), 661–685. https://doi.org/10.1111/j.1469-185X.2011.00216.x

Tsutsui, M. H., Kobayashi, K., & Miyashita, T. (2018). Temporal trends in arthropod abundances after the transition to organic farming in paddy fields. PLOS ONE, 13(1), e0190946. https://doi.org/10.1371/journal.pone.0190946

Tuck, S. L., Winqvist, C., Mota, F., Ahnström, J., Turnbull, L. A., & Bengtsson, J. (2014). Land-use intensity and the effects of organic farming on biodiversity: A hierarchical meta-analysis. Journal of Applied Ecology, 51(3), 746–755. https://doi.org/10.1111/1365-2664.12219

Tylianakis, J. M., Tscharntke, T., & Klein, A.-M. (2006). DIVERSITY, ECOSYSTEM FUNCTION, AND STABILITY OF PARASITOID – HOST INTERACTIONS ACROSS A TROPICAL HABITAT GRADIENT. Ecology, 87(12), 3047–3057. https://doi.org/10.1890/0012-9658(2006)87[3047:DEFASO]2.0.CO;2

Veres, A., Petit, S., Conord, C., & Lavigne, C. (2013). Does landscape composition affect pest abundance and their control by natural enemies ? A review. Landscape ecology and biodiversity in agricultural landscapes, 166, 110–117. https://doi.org/10.1016/j.agee.2011.05.027

Vinatier, F., Gosme, M., & Valantin-Morison, M. (2012). A tool for testing integrated pest management strategies on a tritrophic system involving pollen beetle, its parasitoid and oilseed rape at the landscape scale. Landscape Ecology, 27(10), 1421–1433. https://doi.org/10.1007/s10980-012-9795-3

Zamberletti, P., Papaïx, J., Gabriel, E., & Opitz, T. (2022). Understanding complex spatial dynamics from mechanistic models through spatio-temporal point processes. Ecography, 2022(5), e05956. https://doi.org/10.1111/ecog.05956

Zamberletti, P., Sabir, K., Opitz, T., Bonnefon, O., Gabriel, E., & Papaïx, J. (2021). More pests but less pesticide applications: Ambivalent effect of landscape complexity on conservation biological control. PLoS Computational Biology, 17(11), e1009559. https://doi.org/10.1371/journal.pcbi.1009559

Zollet, S., & Maharjan, K. L. (2021). Overcoming the Barriers to Entry of Newcomer Sustainable Farmers: Insights from the Emergence of Organic Clusters in Japan. Sustainability, 13, 866.

